# Development of AI-assisted microscopy frameworks through realistic simulation in pySTED

**DOI:** 10.1101/2024.03.25.586697

**Authors:** Anthony Bilodeau, Albert Michaud-Gagnon, Julia Chabbert, Benoit Turcotte, Jörn Heine, Audrey Durand, Flavie Lavoie-Cardinal

## Abstract

The integration of artificial intelligence (AI) into microscopy systems significantly enhances performance, optimizing both the image acquisition and analysis phases. Development of AI-assisted super-resolution microscopy is often limited by the access to large biological datasets, as well as by the difficulties to benchmark and compare approaches on heterogeneous samples. We demonstrate the benefits of a realistic STED simulation platform, `pySTED`, for the development and deployment of AI-strategies for super-resolution microscopy. The simulation environment provided by `pySTED` allows the augmentation of data for the training of deep neural networks, the development of online optimization strategies, and the training of reinforcement learning models, that can be deployed successfully on a real microscope.

## 1 Introduction

Super-resolution microscopy has played a pivotal role in life sciences by allowing the investigation of the nano-organization of biological samples to a few tens of nanometers [1]. STimulated Emission Depletion (STED) [2], a point scanning based super-resolution microscopy fluorescence modality, routinely allows resolution down to 30-80 nm to be reached in fixed and live samples [1]. One drawback of STED microscopy is the photobleaching of the fluorophores associated with the increased light exposure at the sample [1, 3, 4]. Photobleaching results in a decrease in fluorescence, limiting the ability to capture multiple consecutive images of a particular area and may also increase phototoxicity in living samples [4, 5]. In an imaging experiment, photobleaching and phototoxicity need to be minimized by careful modulation of the imaging parameters [5, 6] or by adopting smart-scanning schemes [7–9]. Integration of AI-assisted smart-modules to bioimaging acquisition protocols has been proposed to guide and control microscopy experiments [6, 7, 10, 11]. However, Machine Learning (ML) and Deep Learning (DL) algorithms generally require a large amount of annotated data to be trained, which can be difficult to obtain when working with biological samples. Diversity in curated training datasets also enhances the model’s robustness [12, 13]. While large annotated datasets of diffraction-limited optical microscopy have been published in recent years [14, 15], access to such datasets for super-resolution microscopy is still limited, in part due to the complexity of data acquisition and annotation as well as a limited access to imaging resources. Similarly, the development of reinforcement learning (RL) methods adapted to the control of complex systems on a wide variety of tasks in games, robotics, or even in microscopy imaging, are strongly dependent on the availability of large training datasets, generally relying on the development of accessible, realistic, and modular simulation environments [11, 16–20] To circumvent this limitation, simulation strategies have been employed for high-end microscopy techniques. For instance, in Fluorescence Lifetime Imaging Microscopy (FLIM), it is common practice to use simulation software to generate synthetic measurements to train ML/DL models [21]. The models can be completely trained in simulation or with few real measurements. Researchers in Single Molecule Localization Microscopy (SMLM) have also adopted simulation tools in their image analysis pipelines to benchmark their algorithms [22–24]. Nehme *et al*. [25] could train a DL model with simulated ground truth detections and few experimental images which was then deployed on real images. In STED microscopy, simulation software are also available. However, they are limited to theoretical models of the point spread function (PSF) [26, 27] or effective PSF (E-PSF) [8, 28], without reproducing realistic experimental settings influencing the design of STED acquistions (e.g. photobleaching, structures of interest, scanning schemes). This limits the generation of simulated STED datasets and associated training of ML/DL models for smart STED microscopy modules.

We created a simulation platform, `pySTED`, that emulates an *in-silico* STED microscope with the aim to assist the development of AI methods. `pySTED` is founded on theoretical and empirically validated models that encompass the generation of the E-PSF in STED microscopy, as well as a photobleaching model [3, 19, 26, 29]. Additionally, it implements realistic point-scanning dynamics in the simulation process, allowing adaptive scanning schemes and non-uniform photobleaching effects to be mimicked. Realistic samples are simulated in `pySTED` by using a DL model that predicts the underlying structure (datamaps) of real images.

`pySTED` can benefit the STED and machine learning communities by facilitating the development and deployment of AI-assisted super-resolution microscopy approaches (Extended Fig. 1). It is implemented in a *Google Colab* notebook to help trainees develop their intuition regarding STED microscopy on a simulated system (Extended Fig. 1i). We demonstrate how the performance of a DL model trained on a semantic segmentation task of nanostructures can be increased using synthetic images from `pySTED` (Extended Fig. 1ii). A second experiment shows how our simulation environment can be leveraged to thoroughly validate the development of AI methods and challenge their robustness before deploying them in a real-life scenario (Extended Fig. 1iii). Lastly, we show that `pySTED`enables the training of a RL agent that can learn by interacting with the realistic STED environment, which would not be possible on a real system due to data constraints [30]. The resulting trained agent can be deployed in real experimental conditions to resolve nanostructures and recover biologically relevant features by bridging the reality gap (Extended Fig. 1iv).

**Figure 1:**
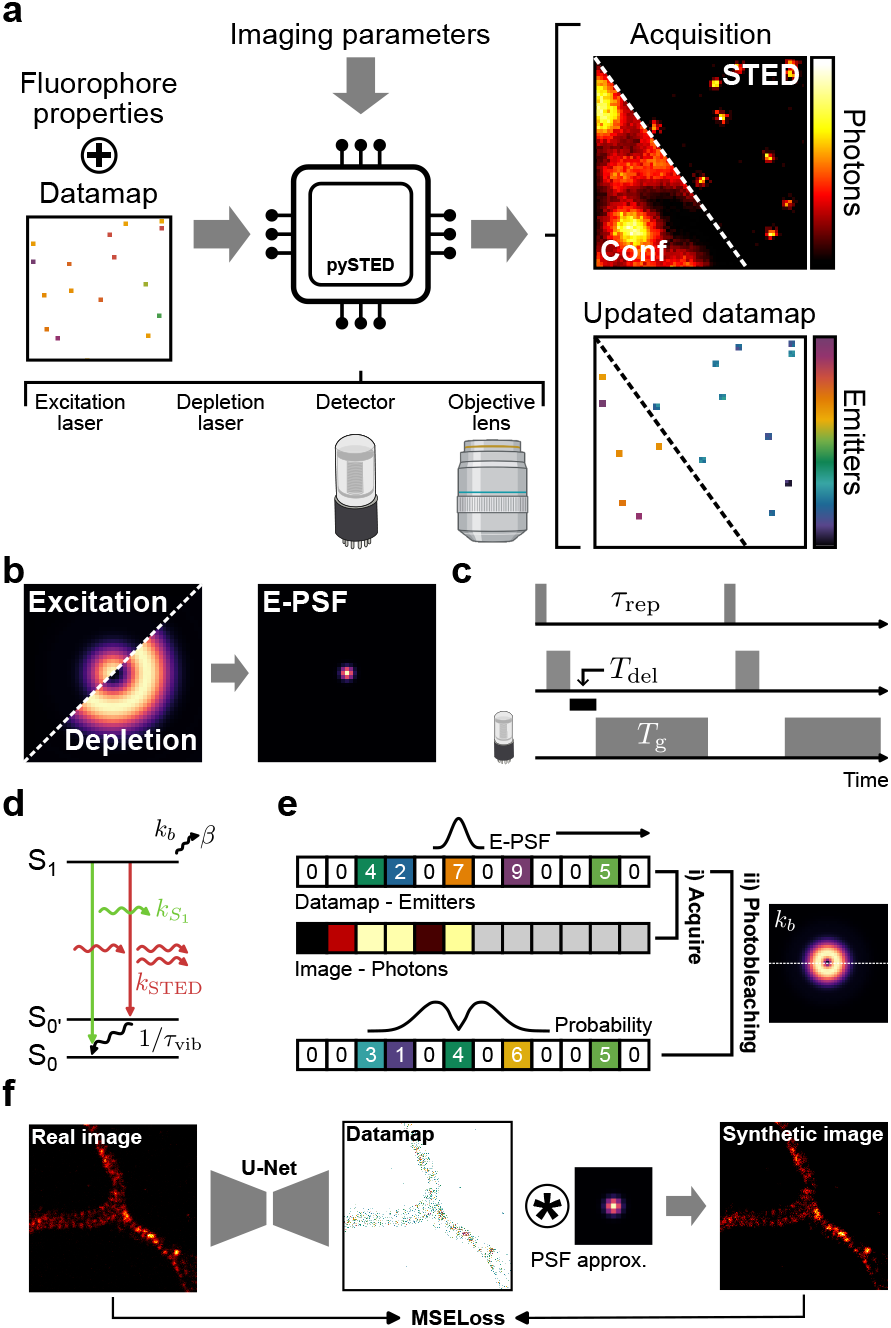
`pySTED` simulation platform. a) Schematic of the `pySTED` microscopy simulation platform. The user specifies the fluorophore properties (e.g. brightness and photobleaching) and the positions of the emitters in the datamap. A simulation is built from several components (excitation and depletion lasers, detector, and objective lens) that can be configured by the user according to their experimental settings. A low-resolution (Conf) or high-resolution (STED) image of an underlying datamap is simulated using the provided imaging parameters. The number of fluorophores on each pixel in the original datamap is updated according to their photophysical properties and associated photobleaching effects. b) Modulating the excitation with the depletion beam impacts the effective point spread function (E-PSF) of the microscope. The E-PSF is convolved on the datamap to calculate the number of photons. c) A time-gating module is implemented in `pySTED`. The temporal acquisition scheme of the simulation can be modulated by the user. It affects the lasers and the detection unit. The time-gating parameters of the simulation (gating delay: *T*_del_ and gating time: *T*_g_) as well as the repetition rate of the lasers (*τ*_rep_) are presented. A grey box is used to indicate when a component is active. d) A two state Jablonski diagram (ground state: *S*0 and excited state: *S*1) presents the transitions that are included in the fluorescence (spontaneous decay: *kS*1 and stimulated emission decay: *k*_STED_) and photobleaching dynamics (photobleaching rate: *k*_*b*_ and photobleached state: *β*) of `pySTED`. The vibrational relaxation rate (1/*τ*_vib_) affects the effective saturation factor in STED. e) An image acquisition is simulated as a two-step process where for each position in the datamap we do the following : i, Acquire) The convolution of the E-PSF with the number of emitters in the datamap (Datamap - Emitters) is calculated to obtain the signal intensity and is reported in the image (Image - Photons). ii, Photobleaching) The number of emitters at each position in the datamap is updated according to the photobleaching probability (line profile from *β*_*b*_, compare top and bottom line). The same colormaps used in **a** are also employed for both the datamap and image in e and f. f) Realistic datamaps are generated from real images. A U-Net model is trained to predict the underlying structure from a real STED image. Convolving the predicted datamap with the approximated PSF results in a realistic synthetic image. During training the mean squared error loss (MSELoss) is calculated between the real and synthetic image. Once trained, the convolution step can be replaced by `pySTED`.

## 2 Results

### 2.1 STED simulation with pySTED

We have built a realistic, open-sourced^1^, STED simulation platform within the Python environment, namely `pySTED`. `pySTED` breaks down a STED acquisition into its main constituents: wavelength dependent focusing properties of the objective lens, fluorophore excitation and depletion, and fluorescence detection. Each step of the acquisition process corresponds to an independent component of the pipeline and is created with its own parameters (Supplementary Tables 1-4) that users can modify according to their experimental requirements (Figure 1a) [26]. Generating a synthetic image with the `pySTED` simulator requires to provide a map of the emitters in the field of view and to specify the photophysical properties of the fluorophore (Figure 1a and Supplementary Table 5). The map of fluorophores, referred to as datamap, can consist of automatically generated simple patterns (e.g. beads, fibers) or more complex structures generated from real images (Methods). The emission and photobleaching properties of the fluorophores that are implemented in `pySTED` are inspired from previous theoretical and experimental models [3, 29]. As in a real experiment, the datamap is continuously being updated during the simulation process to realistically simulate point-scanning acquisition schemes (Figure 1a-e, Methods).

#### Realistic datamap generation

Datamaps that can reproduce diverse biological structures of interest are required for the development of a simulation platform that enables the generation of realistic-*synthetic* STED images. Combining primary object shapes such as points, fibers, or polygonal structures is efficient and simple for some use-cases but is not sufficient to represent more complex and diverse structures that can be found in real biological samples [22–24, 31]. It is essential to reduce the gap between simulation and reality for microscopist trainees or to train artificial intelligence models on synthetic samples prior to the deployment on real tasks [32, 33].

We sought to generate realistic datamaps by training a DL model to predict the underlying structures from real STED images which can then be used in synthetic `pySTED` acquisition. We chose the U-Net architecture, U-Net_datamap_ as it as been shown to perform well on various microscopy datasets of limited size [34, 35] (Figure 1f). We adapted a previously established approach in which a low-resolution image is mapped to a resolution-enhanced image [36, 37]. Once convolved with an equivalent optical transfer function the resolution-enhanced synthetic image is compared with the original image.

Here, we trained the U-Net_datamap_ on STED images of proteins in cultured hippocampal neurons (Methods, Supplementary Fig. 1, and Supplementary Tab. 7). During the training process, the model aims at predicting the underlying structure (datamap) such that the convolution of the approximated PSF of the STED microscope (full-width at half maximum (FWHM): ∼50 nm, measured from FWHM of real STED images) minimizes the mean quadratic error with the real image (Figure 1f). After training, given a real image, the U-Net_datamap_ generates the underlying structure (Supplementary Fig. 1). From this datamap, a synthetic `pySTED` image can be simulated with different imaging parameters (low or high resolution). Qualitative comparison of the synthetic images acquired in `pySTED` with the real STED images (Supplementary Fig. 1) shows similar super-resolved structures for different neuronal proteins confirming the capability of the U-Net_datamap_ to predict a realistic datamap. We also evaluated the quality of images resulting from datamaps generated with the U-Net_datamap_ or a conventional Richardson-Lucy deconvolution (Methods, Supplementary Fig. 2a). As highlighted in Supplementary Fig. 2b,c, the use of the U-Net_datamap_ instead of Richardson-Lucy deconvolution to generate datamaps in `pySTED` results in improved synthetic images.

**Figure 2:**
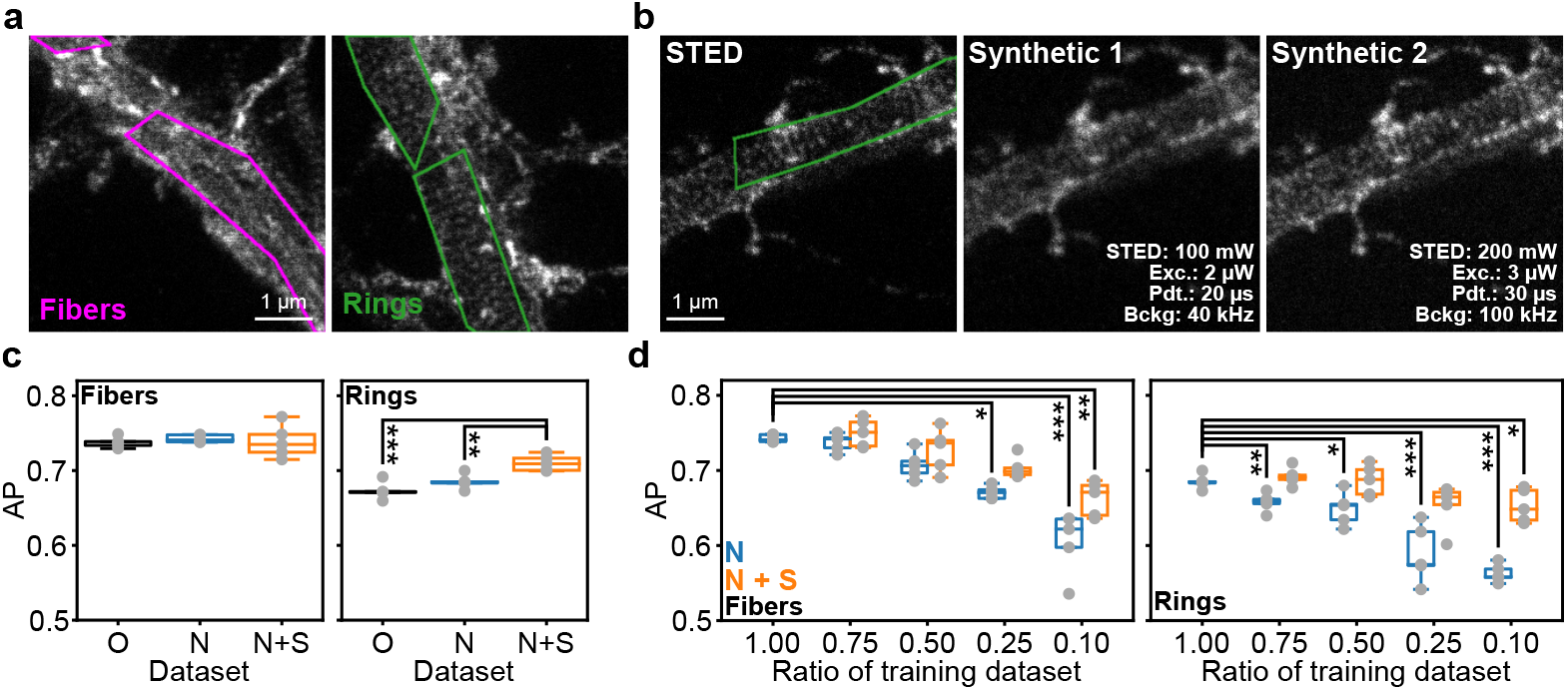
`pySTED` is used to artificially augment the training dataset of a DL model. a) We tackle the segmentation task that was used in Lavoie-Cardinal *et al*. [43] where the annotations consist in polygonal bounding boxes around F-actin fibers (magenta) and rings (green). b) `pySTED` is used to augment the training dataset by generating synthetic versions of a STED image. c) Average Precision (AP) of the model for the segmentation of F-actin fibers (magenta) and rings (green). The model was trained on the original dataset from Lavoie-Cardinal *et al*. [43] (O), and on the same dataset with updated normalization (N) and additionnal synthetic images (N+S). No significant changes in AP are measured for F-actin fibers but a significant increase is measured for N+S over O and N for F-actin rings (*p*-values in Supplementary Fig. 6). d) Images were progressively removed from the dataset (100%: 42 images, 75%: 31 images, 50%: 21 images, 25%: 10 images, and 10%: 4 images). Removing more than 50% of the dataset for fibers negatively impacts the models whereas removing 25% of the dataset negatively impacts the segmentation of rings (N; *p*-values in Supplementary Fig. 6). Adding synthetic images from `pySTED` during training allows 75% of the original training dataset to be removed without affecting the performance for both structures (N + S, *p*-values in Supplementary Fig. 6). Only the significant changes from the complete dataset are highlighted. The complete statistical analysis is provided in Supplementary Fig. 6.

#### Validation of pySTED with a real STED microscope

We characterized the capacities of `pySTED` to simulate realistic fluorophore properties by comparing the synthetic `pySTED` images with real STED microscopy acquisitions. We acquired STED images of the protein bassoon, which had been immunostained with the fluorophore ATTO-647N in dissociated cultured hippocampal neurons. We compared the effect of varying the imaging parameters on the `pySTED` simulation environment and on the real microscope (Supplementary Fig. 3-5). For `pySTED` we used the photophysical properties of the fluorophore ATTO-647N from the literature (Supplementary Tab. 5) [3, 38]. The photobleaching constants (k_1_ and b) were estimated from the experimental data by using a least-squares fitting method (Methods). Synthetic datamaps were generated with the U-Net_datamap_ to facilitate the comparison between simulation and reality.

**Figure 3:**
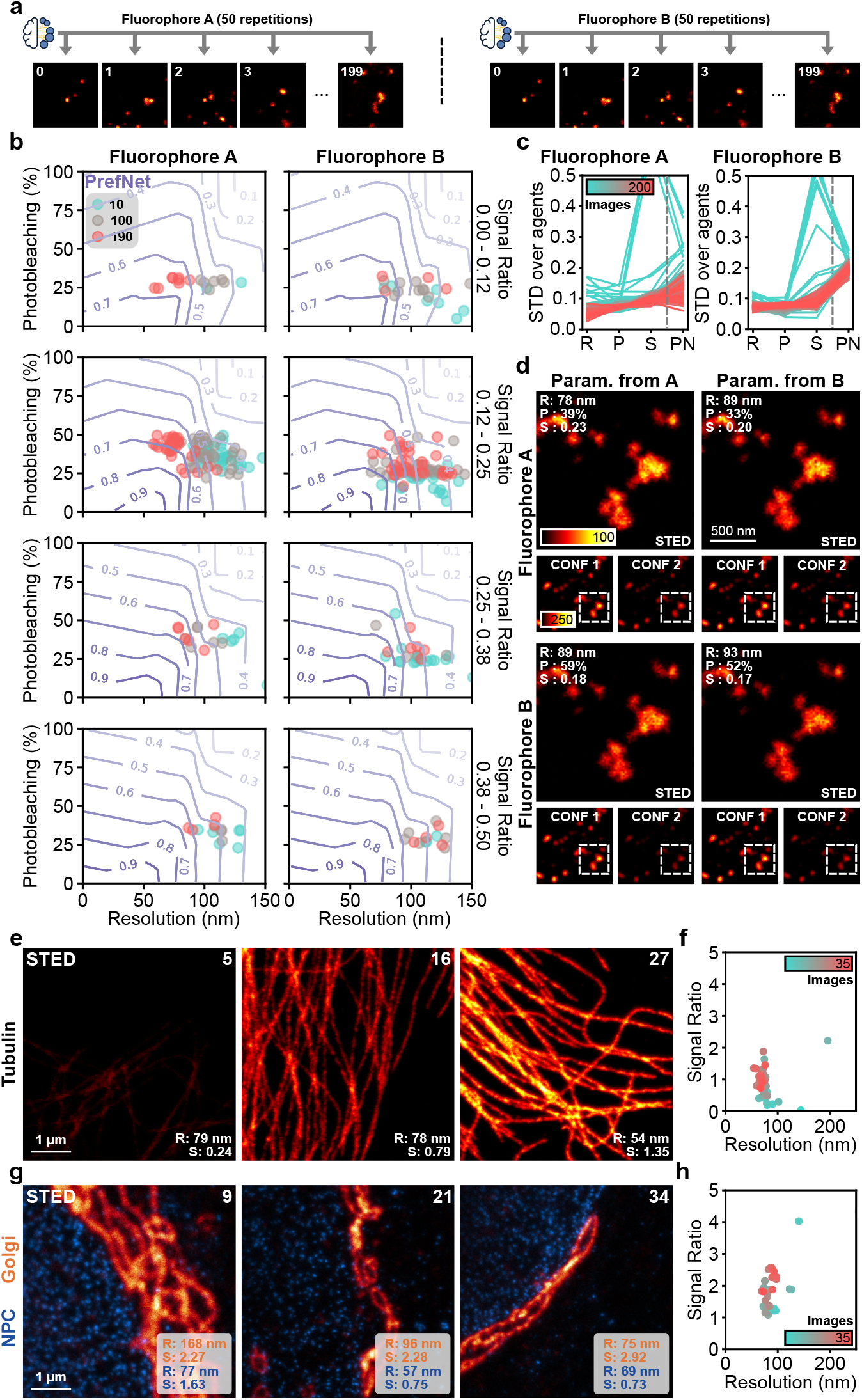
Validation of AI-assisted algorithms with `pySTED` for STED microscopy parameter optimization. a) `pySTED` is used to confirm the robustness of a model to the random initialization by repeatedly optimizing (50 repetitions) the imaging parameters on the same sequence of datamaps (200 images). Two fluorophores are considered for demonstration purposes (Supplementary Tab. 9). b) Resulting imaging optimization objectives from LinTSDiag at 3 different timesteps (10 - cyan, 100 - grey, and 190 - red) for 50 independent models which are presented for increasing signal ratio (top to bottom). With time, LinTSDiag acquires images that have a higher preference score for both fluorophores (purple contour lines) and converges into a similar imaging optimization objective space (red points). c) The standard deviation (STD) of the imaging optimization objectives and of the preference scores decreases during the optimization (cyan to red) supporting the convergence of LinTSDiag in a specific region of the imaging optimization objective space for both fluorophores. The dashed line separates the imaging optimization objectives (R: Resolution, P: Photobleaching, and S: Signal ratio) from the preference network (PN). d) Typical `pySTED` simulations on two different fluorophores (top/bottom) using the optimized parameters on fluorophore A (left) or B (right). Parameters that were optimized for fluorophore A (top-left) result in higher photobleaching with maintaining a similar resolution and signal ratio on fluorophore B (bottom-left) compared to parameters that were optimized for fluorophore B (bottom-right). See Supplementary Tab. 9 for imaging parameters. e) Example acquisition of LinTSDiag on a Tubulin in kidney epithelial cells (Vero cells) stained with STAR RED in the beginning (left) and at the end of the optimization (right). f) Over time, LinTSDiag manages to increase both the resolution and the signal ratio of the acquired images (35 images, cyan to red). g) LinTSDiag allows multi-color imaging due to it’s high dimensional parameter space capability. LinTSDiag optimizes the averaged resolution and signal ratio from both channels in dual-color images acquired of Golgi (STAR ORANGE) and Nuclear Pore Complex (STAR RED) in Vero cells. h) LinTSDiag can maximize the signal ratio in the images while maintaining the resolution of the images (35 images, cyan to red).

We first compared how the imaging parameters on the real microscopes and in the `pySTED` simulations (pixel dwelltime, excitation and depletion powers) influenced the image properties by measuring the resolution [39] and the signal ratio [6] (Methods and Supplementary Fig. 3a). As expected, modulating the STED laser power influences the spatial resolution in real experiments and in `pySTED` simulations. Examples of acquired and synthetic images are displayed in Supplementary Fig. 3b for visual comparison with different parameter combinations (Supplementary Fig. 3a). The impact of the imaging parameters in the resolution and signal ratio metrics in `pySTED` agree with the measurements that were performed on a real microscope. The small deviations can be explained by the variability that is typically observed in the determination of absolute values of fluorophore properties [40].

Next, we validated the photobleaching model that is implemented within `pySTED`. We calculated the photobleaching by comparing the fluorescence signal in a low-resolution image acquired before (CONF1) and after (CONF2) the high-resolution acquisition [6] (Methods). For the pixel dwelltime and the excitation power we measured similar trends between real and synthetic image acquisitions (Supplementary Fig. 4a). For a confocal acquisition, the photobleaching in `pySTED` is assumed to be 0 (Supplementary Fig. 4a) as it is generally negligible in a real confocal acquisition. Considering the flexibility of `pySTED`, different photobleaching dynamics specifically tailored for any particular experiment can be implemented and added in the simulation platform. Examples of sequential acquisition (10 images) are presented in Supplementary Fig. 4b to demonstrate the effect of the imaging parameters on the photobleaching of the sample. `pySTED` also integrates background effects that can influence the quality of the acquired images as in real experiments [41, 42] (Supplementary Fig. 4c,d).

**Figure 4:**
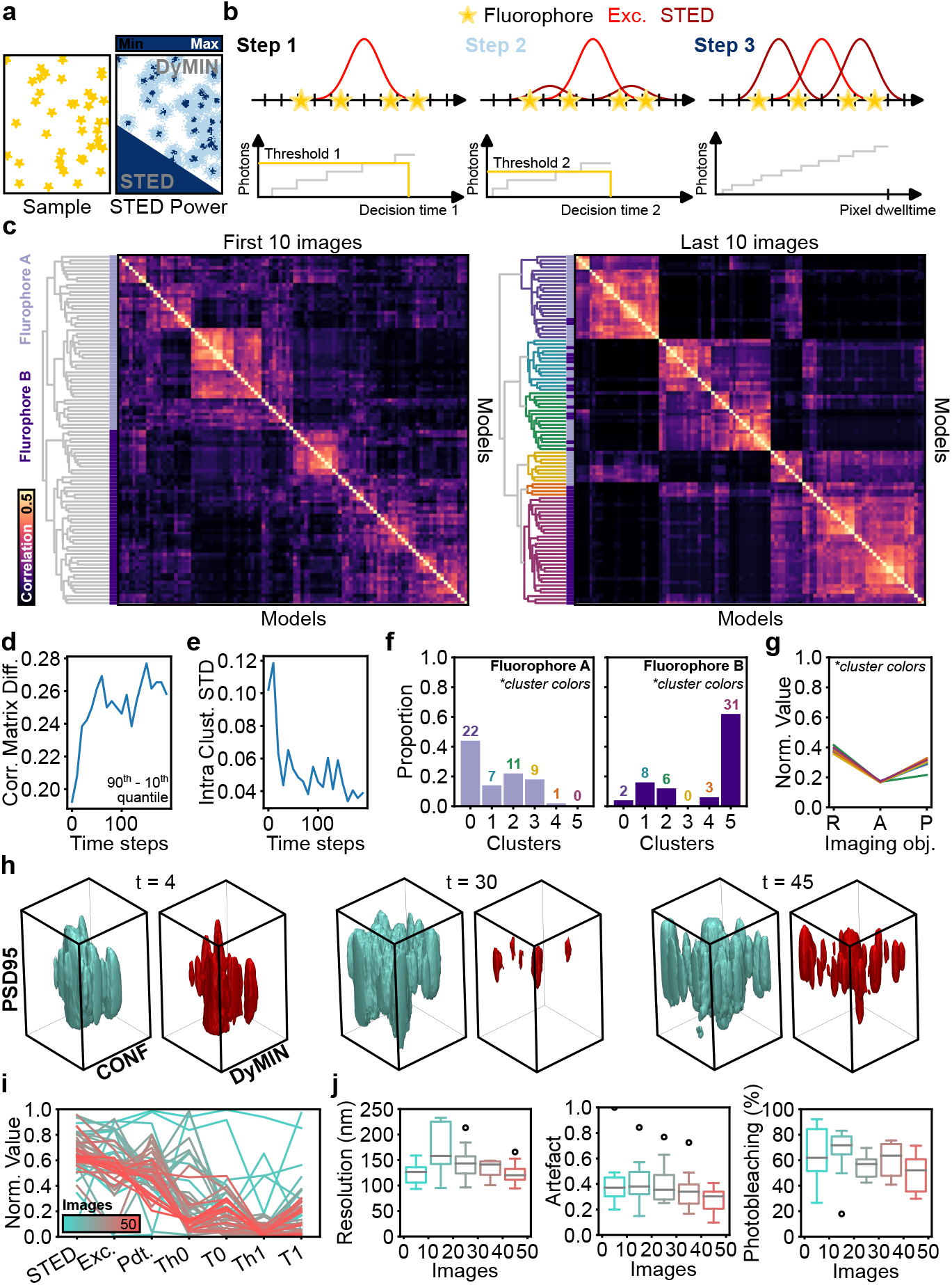
Validation of contextual-bandit algorithms with `pySTED` in a high-dimensional parameter space. a) DyMIN microscopy uses thresholds to turn off the high-intensity STED laser when no structures are within the vicinity of the donut beam (white regions). Thus limiting the light doses at the sample compared to conventional STED. b) Typically DyMIN uses a 3 step process at each pixel. In the first step, only the excitation (Exc.) laser is used and the signal is measured. If the measured signal is higher than the predefined threshold (Threshold 1) after the decision time (Decision Time 1) then the depletion power (STED) is slightly increased and the signal is measured again (Threshold 2 and Decision time 2). Otherwise the acquisition is stopped until the next pixel. The final step (Step 3) consists in a normal STED acquisition. c) `pySTED` was used to characterize LinTSDiag models that can simultaneously optimize 7 parameters (STED and excitation powers, pixel dwelltime, threshold 1 & 2, and decision time 1 & 2) with prior information about the task (confocal image). The convergence of the models to similar parameter combinations is evaluated by measuring the correlation in the action selection (50 models) over time (See Supplementary Fig. 8). Clustering of the correlation matrix reveals clusters of policies that are better defined later in the optimization process (right dendrogram, color-coded). The shades of purple on the left of the correlation matrix represent two different fluorophores (light: A, dark: B). d) The difference between the 90^th^ and 10^th^ quantile of the correlation matrix increases with time implying better defined clusters of policies. e) The intra cluster standard deviation (STD) of the parameter selection decreases during the optimization showing that the policy of the models converges in all defined clusters. f) The proportion of models per cluster for fluorophore A or B (light and dark respectively) shows that there are different modes of attraction in the parameter space for fluorophores with distinct photophysical properties (color-code from c). g) While models converged in different regions of the parameter space, the measured imaging optimization objectives (R: Resolution, A: Artefact, P: Photobleaching) are similar for each cluster (color-code from c). h) Example acquisition with LinTSDiag optimization on a real acquisition task for DyMIN3D of the synaptic protein PSD95 in cultured hippocampal neurons. The volume size is 2.88 µm *×* 2.88 µm *×* 2 µm. Confocal (left) and DyMIN (right) acquisitions are displayed. i) A convergence of the parameter selection in the 7-parameter space is observed (cyan to red, STED: STED power, Exc.: Excitation power, Pdt.: Pixel dwelltime, Th1-2: First and second DyMIN threshold, and T1-2: First and second DyMIN decision time). j) LinTSDiag optimization reduces the variability of all imaging optimization objectives during the optimization (50 images). Boxplot shows the distribution in bins of 10 images.

### 2.2 pySTED as a development platform for AI-assisted microscopy

#### 2.2.1 Dataset augmentation for training deep learning models

DL models are powerful tools to rapidly and efficiently analyse large databanks of images and perform various tasks such as cell segmentation [35, 44]. When no pretrained models are readily available online to solve the task [45], finetuning or training a DL model from scratch requires the tedious process of annotating a dataset. We herein aim to reduce the required number of distinct images for training by using `pySTED` as an additional data augmentation step. As a benchmark, we used the F-actin segmentation task from Lavoie-Cardinal *et al*. [43], where the goal is to segment dendritic F-actin fibers or rings using a small dataset (42 images) of STED images (Figure 2a, Methods). `pySTED` was used first as a form of data augmentation to increase the number of images in the training dataset without requiring new annotations. Using U-Net_datamap_ we generated F-actin datamaps and a series of synthetic images in `pySTED` with various simulation parameters (Figure 2b, Supplementary Tab. 8).

We compared the segmentation performance by using the average precision (AP, Methods) of a DL model trained on the original dataset (*O* [43]) or with different image normalization and increased data augmentation (*N*). The segmentation performance was not impacted by increasing the amount of data augmentation (O vs. N, Figure 2c). Adding synthetic images from `pySTED` (N+S) into the training dataset to improve the diversity of the dataset significantly increases the performance of F-actin rings segmentation compared to *O* and *N* and maintains the performance for the F-actin fibers segmentation (Figure 2c). In biological experiments, where each image is costly to acquire, reducing the size of the training dataset results in a higher number of images for the post-hoc analysis. Hence, we sought to measure the impact of reducing the number of real images in the training dataset by training on subsets of images that are augmented using `pySTED` (Supplementary Fig. 7). We measure a significant decrease of the AP for F-actin fibers when the model is trained on less than 50% of the images. Removing 25% of the dataset negatively impacts the segmentation performance of F-actin rings (Figure 2d, *p*-values in Supplementary Fig. 6). However, adding synthetic images from `pySTED` during training allows the segmentation performance of the model to be maintained by training with only 25% of the original dataset (Figure 2d, p-values in Supplementary Fig. 6).

#### 2.2.2 Validation of AI methods

Benchmarking AI models for automating microscopy tasks on biological samples is challenging due to biological variability and the difficulty of comparing imaging strategies on the same region of interest [6, 22, 46]. Assessing and comparing AI models requires multiple attempts in similar, yet different experimental conditions to limit the impact of biological variability. This inevitably increases the number of biological samples and the time required to develop robust AI-assisted adaptive microscopy strategies that can be deployed on a variety of samples and imaging conditions. `pySTED` allows the simulation of multiple versions of the same images as if the structure had been imaged with different experimental settings. We herein showcase the capability of `pySTED` in thoroughly validating ML approaches for the optimization of STED imaging parameters in a simulated controlled environment, enabling more robust performance assessments and comparisons.

We first demonstrate how `pySTED` can be used to characterize the performance of a multi-armed bandit optimization framework that uses Thompson Sampling (TS) for exploration, Kernel-TS. The application of Kernel-TS for the optimization of STED imaging parameters was demonstrated previously, but comparison between different experiments was challenging due to local variations in the size, brightness, and photostability of the fluorescently tagged neuronal structures [6]. Using synthetic images generated with `pySTED` allows the performance of Kernel-TS to be evaluated on the same image sequence (50 repetitions, Methods) and with controlled photophysical properties of fluorophores (Extended Fig. 2 and Supplementary Tab. 14). For experimental settings such as multi-channel imaging or adaptive scanning, Kernel-TS is limited by the number of parameters that can be simultaneously optimized (∼4) in an online setting [6]. We thus turned to a neural network implementation of Thompson Sampling which was recently developed to solve the multi-armed bandit framework, LinTSDiag [47].

Using `pySTED` we could characterize the performance of LinTSDiag on a microscopy optimization task on synthetic images without requiring real biological samples. As described above, LinTSDiag was trained on the same sequence (50 repetitions, Methods) using two different fluorophores (Figure 3a and Supplementary Tab. 14). In a simple 3-parameters optimization setting, LinTSDiag allows a robust optimization of the signal ratio, photobleaching and spatial resolution for fluorophores with distinct photophysical properties (Figure 3b). We evaluate the performance of LinTSDiag using the preference score, which is obtained from a network that was trained to predict the preferences of an expert in the imaging optimization objective space (PrefNet, see Methods) [6]. The convergence of the agent in the imaging optimization objective space is supported by the smaller standard deviation measured in the last iterations of the imaging session (red lines, Figure 3c). `pySTED` enables the comparison of the optimized parameters for different fluorophores on the same datamap. This experiment confirms that optimal parameters vary depending on the photophysical properties (Figure 3d).

LinTSDiag was then deployed on a real microscopy system to simultaneously optimize 4 parameters (Excitation power, STED power, pixel dwelltime, and linesteps) for the imaging of Tubulin stained with STAR RED in kidney epithelial cells (Vero cell line). The model was able to optimize the imaging optimization objectives, improving the resolution and signal ratio, while maintaining a low level of photobleaching over the course of the optimization (Figure 3e,f and Supplementary Tab. 14). Then we sought to increase the number of parameters by tackling a dual-color imaging scheme (6 parameters, Excitation power, STED power, and linesteps for both channels) for STED imaging of Golgi stained with STAR-ORANGE and nuclear pore complex (NPC) stained with STAR RED in Vero cells (Figure 3g,h and Supplementary Tab. 14). The optimization framework allows 4 imaging optimization objectives to be simultaneously optimized (*e*.*g*. resolution and signal ratio for both colors). As the visual selection of the trade-off in a 4-dimensional space is challenging for the user in an online setting, we decided to optimize the combined resolution and signal ratio of both fluorophores (average of the imaging optimization objectives), allowing the users to indicate their preference in a two-dimensional optimization objective space. Online 6-parameters optimization of LinTSDiag increases the signal ratio while maintaining a good image resolution for both imaging channels (Figure 3h) enabling to resolve both structures with sub-100 nm resolution.

Next, we developed a model that leverages prior information (context) to solve a task with a high-dimensional action space. This is the case for DyMIN microscopy which requires parameter selection to be adapted, in particular multiple illumination thresholds, to the current region of interest [8] (Figure 4a,b). We previously showed that contextual-bandit algorithms can use the confocal image as a context to improve DyMIN thresholds optimization in a two parameters setting [48]. In this work we aim to increase the number of parameters (7 parameters) that can be simultaneously optimized and validate the robustness of LinTSDiag [47] (Figure 4b). We repeatedly trained LinTSDiag on the same datamap sequence using the confocal image as prior information (50 repetitions). The parameter selection was compared by measuring whether the action selection correlated over time between the models (Figure 4c, Supplementary Fig. 8, Supplementary Tab. 14, and Methods). For instance, the correlation matrix from the last 10 images shows clusters of similar parameters that are better defined than for the first 10 images (Figure 4c). This is confirmed by the 90^th^ and 10^th^ quantile difference in the correlation matrix which rapidly increases with time (Figure 4d). As expected with clustered policies, the average standard deviation of the action selection for each cluster reduces over time implying similar parameter selection by the models (Figure 4e). We also assessed whether the models would adapt their policies to different fluorophores (light/dark purple, Figure 4c,f). As shown in Figure 4f, there are specific policies for each fluorophore (*e*.*g*. fluorophore A: 0, 3; fluorophore B: 5) demonstrating the capability of the models in adapting their parameter selections to the experimental condition. While the policy of the models are different, the measured imaging optimization objectives are similar for all clusters (Figure 4g) which suggests that different policies can solve this task unveiling the delicate intricacies of DyMIN microscopy. More importantly this shows that the model can learn one of the many possible solutions to optimize the imaging task.

The LinTSDiag optimization strategy was deployed in a real life experiment for the 7 parameter optimization of DyMIN3D imaging of the post-synaptic protein PSD95 in dissociated primary hippocampal neurons stained with STAR-635P. Early in the optimization, the selected parameters produced images with poor resolution or missing structures (artefacts) (Figure 4h and Supplementary Tab. 14). The final images were of higher quality (right, Figure 4h) with fewer artefacts and high resolution. The parameter selection of the model converged in a region of the parameter space that could improve all imaging optimization objectives over the course of optimization (Figure 4i,j). Parameters optimized with LinTSDiag allowed a significant improvement of DyMIN3D imaging of PSD95 compared to conventional 3D STED imaging (Supplementary Fig. 9). `pySTED` allowed us to validate the robustness of the model in a simulated environment prior to its deployment in a real experimental setting. This should benefit the ML community by allowing the validation of new online ML optimization algorithms on realistic tasks.

### 2.3 Learning through interactions with the system

Online optimization strategies such as Kernel-TS and LinTSDiag were trained from scratch on a new sample, implying a learning phase in which only a fraction of the images will meet minimal image quality requirements. For costly biological samples, there is a need to deploy algorithms that can make decisions based on the environment with a reduced initial exploration phase. Control tasks and sequential planning are particularly well suited for a RL framework where an agent (e.g. replacing the microscopist) learns to make decisions by interacting with the environment (e.g. select imaging parameters on a microscope) with the aim of maximizing a reward signal (e.g. light exposure, signal ratio, resolution) over the course of an episode (e.g. imaging session) [49]. Deep RL agents are (unfortunately) famously data-intensive, sometimes requiring millions of examples to learn a single task [17, 30]. This makes them less attractive to be trained on real-world tasks where each sample can be laborious to obtain (e.g. biological samples) or when unsuitable actions can lead to permanent damage (e.g. overexposition of the photon detector). Simulation platforms are thus essential in RL to provide environments in which an agent can be trained at low cost to then be deployed in a real-life scenario [50], which is referred to as simulation to reality (Sim2Real) in robotics. While Sim2Real is widely studied in robotics and autonomous driving, its success for new fields of application is generally dependant on the gap between simulation and reality [51].

Here, `pySTED` is used as a simulation software to train RL agents. We implemented `pySTED` in an OpenAI Gym environment (`gym-STED`) to facilitate the deployment and development of RL strategies for STED microscopy [19, 52]. To highlight the potential of `gym-STED` to train a RL agent, we crafted the task of resolving nanostructures in simulated datamaps of various neuronal structures (Figure 5a). In `gym-STED` an episode unfolds as follows. At each timestep the agent observes the state of the sample: a visual input (image) and the current history (Methods, Figure 5b). The agent then performs an action (adjusting pixel dwelltime, excitation and depletion powers), receives a reward based on the imaging optimization objectives and transitions into the next state. A single value reward is calculated using a preference network that was trained to rank the imaging optimization objectives (resolution, photobleaching and signal ratio) according to expert preferences [6] (Methods). A negative reward is obtained when the selected parameters lead to a high photon count that would be detrimental to the detector in real experimental settings (*e*.*g*. non-linear detection of photon counts). This sequence is repeated until the end of the episode, 30 timesteps. In each episode, the goal of the agent is to balance between detecting the current sample configuration and acquiring high-quality images to maximize it’s reward (Figure 5a). We trained a proximal policy optimization (PPO) [53] agent and evaluate its performance on diverse fluorophores (Methods). Domain randomization is used heavily within the simulation platform to cover a wide variety of fluorophores and structures and thus increase the generalization properties of the agent [54]. In Figure 5c-f (Supplementary Tab. 15), we report the performance of the agent on a fluorophore with simulated photophysical properties that would result in high brightness (high signal ratio) and high photostability (low photobleaching) in real experiments. The results of the agent on other simulated fluorophore properties are reported in supplementary material (Supplementary Tab. 10 and Supplementary Fig. 10). Over the course of training, the agent adapts its policy to optimize the imaging optimization objectives (100k and 12M training steps, Figure 5c). As expected from RL training, the reward of an agent during an episode is greater at the end of training compared to the beginning (red vs. cyan, Figure 5d). When evaluated on a new sequence, the agent trained over 12M steps rapidly adapts its parameter selection during the episode to acquire images with high resolution and signal ratio, while minimizing photobleaching (Figure 5e,f). The agent shows a similar behavior for various simulated fluorophores (Supplementary Fig. 10). We compared the number of good images acquired by the RL agent with that of bandit optimization for the first 30 images of the optimization. In similar experimental conditions, with the same fluorophore and parameter search space, the average number of good images were (18±3) and (5±3) for the RL agent and bandit respectively (50 repetitions). This almost four-fold increase in the number of high quality images, highlights the improved efficiency of the RL agent at suggesting optimal imaging parameters.

**Figure 5:**
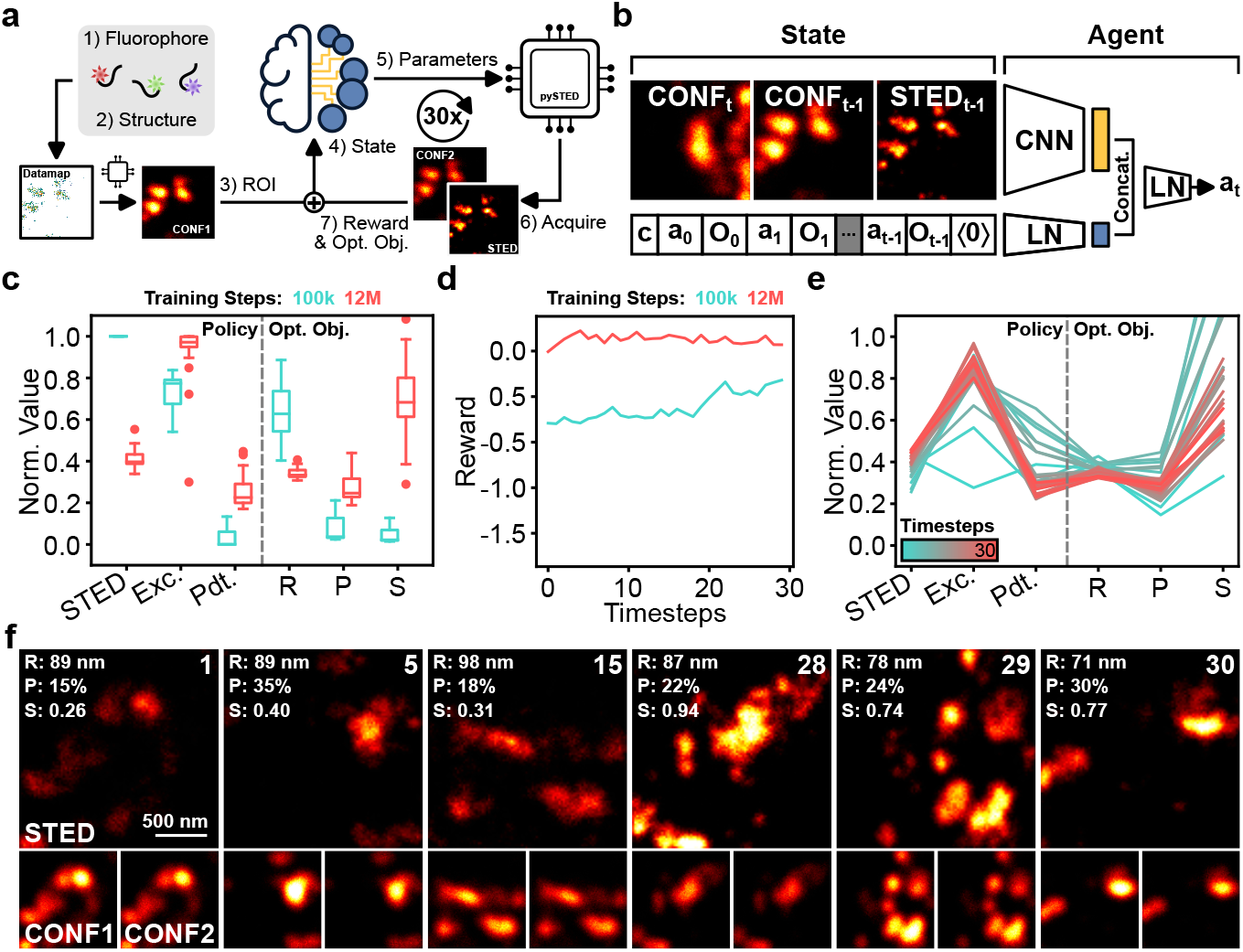
A RL agent is trained to optimize the STED imaging parameters in simulation with `pySTED`. a) Schematic of the RL training loop in simulation. Each episode starts by sampling a set of photophysical properties representing a fluorophore (1) and the selection of a structural protein from the databank (2). At each timestep a region of interest (ROI) is selected: a datamap is created and a confocal image is generated with `pySTED` (3). The confocal image is used in the state of the agent (4) which then selects an action, *i*.*e*. the next imaging parameters (5). A STED image and a second confocal image are generated in `pySTED` (6). The imaging optimization objectives and the reward are calculated (7). On the next timestep, the agent sees a new ROI, the previously simulated images and the history of the episode. b) The state of the agent includes a visual input (the images) and the history. The visual input of the agent is the current confocal (CONF*t*) and the previous confocal/STED images (CONF_*t-*1_ and STED_*t-*1_). The state of the agent also incorporates the laser excitation power at which the confocal image was acquired (*c*), the history of selected actions (*a*_*t*_) and the calculated imaging optimization objectives (*O*_*t*_). The history vector is zero-padded to a fixed length (⟨0⟩). The agent encodes the visual information using a convolutional neural network (CNN) and the history using a fully connected linear layer (LN). Both encoding are concatenated and fed to a LN model which predicts the next action. c) Evolution of the policy (left, STED: STED power, Exc.: Excitation power, Pdt.: Pixel dwelltime) and imaging optimization objectives (right, R: Resolution, P: Photobleaching, S: Signal ratio) for a fluorophore with high-signal and low-photobleaching properties during training at the beginning (cyan, 100k timesteps) and at the end (red, 12M timesteps) of the training process. A boxplot shows the distribution of the average value from the last 10 images of an episode (30 repetitions). d) Evolution of the reward during an episode at the beginning (cyan, 100k timesteps) and at the end of training (red, 12M timesteps) for the same fluorophore properties as in c). e) Evolution of the policy (left) and imaging optimization objectives (right) after training (12M timesteps) during an episode for a fluorophore with the same photophysical properties as in c). f) Typical examples of images acquired during an episode. The image index is shown in the top right corner and the calculated imaging optimization objectives in the top left corner. The STED image and second confocal (CONF2) image are normalized to their respective first confocal (CONF1) images.

**Figure 6:**
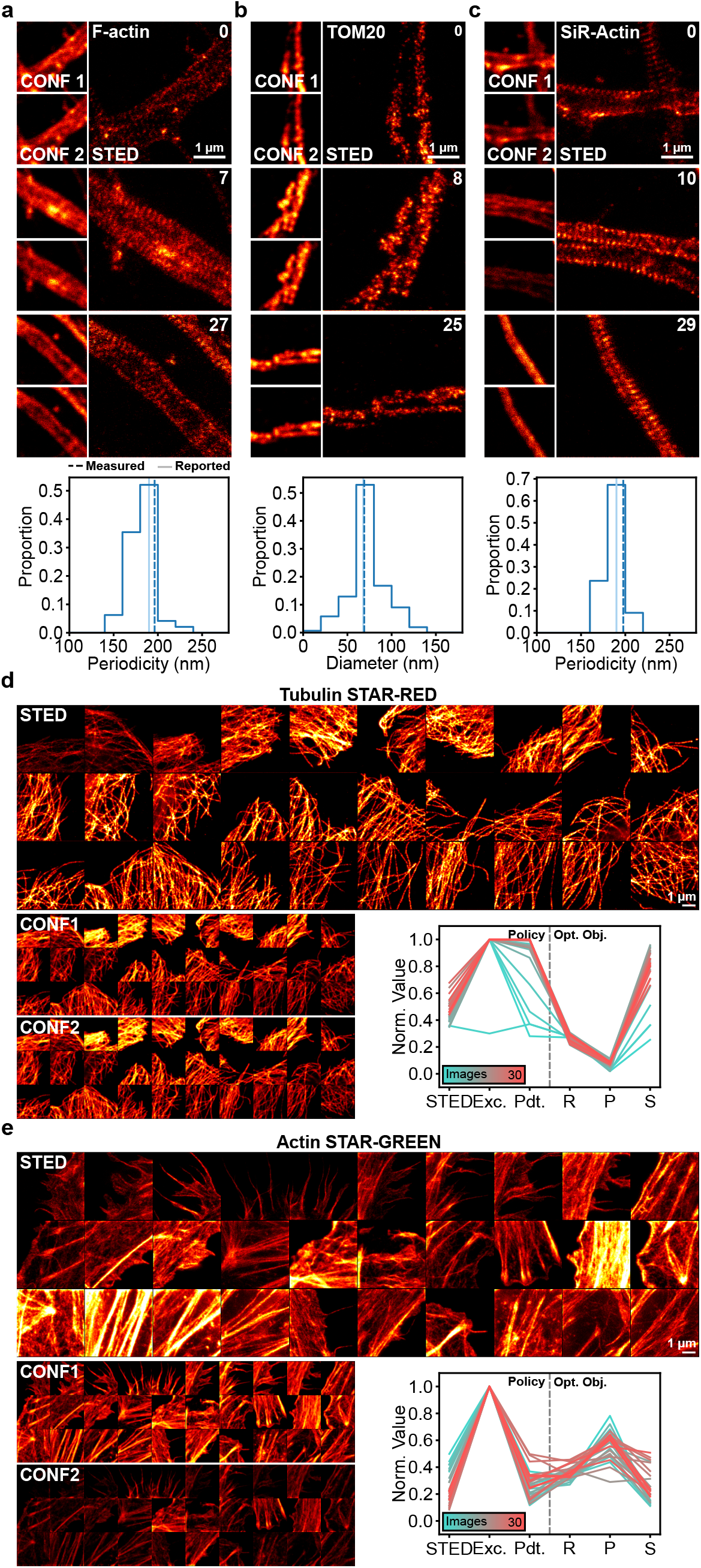
Bridging the reality gap between simulation and reality in RL by pretraining with `pySTED`. For all real microscopy experiments, the deployed agent was trained over 12M steps in simulation. The agent was deployed on a real STED microscope for the imaging of diverse proteins in dissociated neuronal cultures and cultivated Vero cells. a) *Top*: Simulated images of F-actin in fixed neurons were used during the training process. Deploying the RL agent to acquire an image of this *in distribution* structure in a real experiment allows the periodic lattice of F-actin tagged with Phalloidin-STAR635 to be revealed in all acquired images. *Bottom*: Structural parameters are extracted from the acquired images (the dashed vertical line represents the median of the distribution) and compared to the values that were previously reported in the literature (solid vertical line). The agent has learned to adjust the imaging parameters to resolve the 190 nm periodicity of the F-actin periodic lattice [55, 56]). b) *Top*: The trained agent is tested on the protein TOM20, a structure that was never seen during training (*out of distribution*). The nano-organization of TOM20 is revealed in all acquired images. *Bottom*: The measured average cluster diameter of TOM20 concords with the averaged reported values from Wurm *et al*. [57]. c) *Top*: Live-cell imaging of SiR-Actin shows the capacity of the model in adapting to different experimental conditions (*out of distribution*). *Bottom*: The periodicity of the F-actin lattice is measured from each acquired images and compared with the literature. See Material and Methods for the quantification. The STED images are normalized to their respective confocal image (CONF1). The second confocal image (CONF2) uses the same colorscale as CONF1 to reveal photobleaching effects. d,e) Images acquired by the RL agent in a real experiment on a different microscope. Tubulin was stained with the STAR-RED fluorophore (d) and Actin was stained with STAR-GREEN (e)in fixed Vero cells. The sequence of acquired images goes from top left to bottom right. The confocal images before (CONF1) and after (CONF2) are presented for photobleaching comparison. The CONF1 image is normalized to the CONF2 image. The STED images are normalized to the 99^th^ percentile of the intensity of the CONF1 image. Images are 5.12 µm *×* 5.12 µm. The evolution of the parameter selection (left; STED: STED power, Exc.: Excitation power, Pdt.: Pixel dwelltime) and imaging optimization objectives (right; R: Resolution, P: Photobleaching, S: Signal ratio) are presented, showing that optimal parameters and optimized objectives for STAR-RED (d) and STAR-GREEN (e) can differ greatly.

Given the capability of the agent in acquiring images for a wide variety of synthetic imaging sequences, we evaluated if the agent could be deployed in a real experimental setting. The experimental conditions chosen for the simulations were based on the parameter range available on the real microscope. Dissociated primary hippocampal neurons were stained for various neuronal proteins (Figure 6, Extended Fig. 3, and Supplementary Tab. 15) and imaged on a STED microscope with the RL agent assistance for parameter choice. First, we evaluated the performance of our approach for *Sim2Real* on *in distribution* images from F-actin and CaMKII-,*β* in fixed neurons. While simulated images of both structures were available within the training environment we wanted to evaluate if the agents could adapt to the real life imaging settings (Supplementary Fig. 1). As shown in Figure 6a and Extended Figure 3, the agent resolves the nano-organization of both proteins (Supplementary Fig. 11). We sought to confirm whether the quality of the images was sufficient to extract biologically relevant features (Methods). For both proteins, the measured quantitative features matched with values previously reported in the literature, enabling the resolution of the 190 nm periodicity of the F-actin lattice in axons, and the size distribution of CaMKII-,*β* nanoclusters [55, 58] (Figure 6a and and Extended Figure 3). Next, we wanted to validate that the agent would adapt it’s parameter selection to structures, fluorophores properties or imaging conditions that were not included in the training set. We first observed that the agent could adapt to a very bright fluorescent signal and adjust the parameters to limit the photon counts on the detector (Extended Fig. 3). The morphology of the imaged PSD95 nano-cluster was in agreement with the values reported by Nair *et al*. [59] (Extended Fig. 3). We deployed the RL-based optimization scheme for the imaging of the mitochondrial protein TOM20 to evaluate the ability of the agent to adapt to *out-of-distribution* structures (Figure 6b). The nano-organization and morphology previously described by Wurm *et al*. [57] of TOM20 in punctate structures is revealed using the provided imaging parameters in all acquired images (Figure 6b and Supplementary Fig. 11). Next, we evaluated the generalizability of the approach to a new imaging context, which is live-cell imaging. We used the optimization strategy for the imaging of the F-actin periodic lattice in living neurons (Figure 6c). The quality of the acquired images are confirmed by the quantitative measurement of the periodicity which matches the previously reported values of 190 nm from the literature [55, 56]. Finally, we verified the generalizability of our approach by deploying our RL-assisted strategy on a new microscope and samples (Figure 6d-e, Extended Fig. 4, and Supplementary Tab. 15). We evaluated the performance of the RL agent for the imaging in fixed Vero cells of tubulin stained with STAR-RED and Actin stained with STAR-GREEN. The agent successfully adapted to the new imaging conditions, rapidly acquiring high quality images, even in challenging photobleaching conditions such as with STED microscopy of the green emitting fluorophore STAR-GREEN. Using the `pySTED` simulation environment we could successfully train RL agents that can be deployed in a variety of real experimental settings to tackle STED imaging parameter optimization tasks. To our knowledge, this is the first application of RL agents to an online image acquisition task in optical microscopy.

## 3 Discussion

We built `pySTED`, an *in-silico* super-resolution STED environment, which can be used to develop and benchmark AI-assisted STED microscopy. Throughout synthetic and real experiments, we have demonstrated that it can be used for the development and benchmarking of AI approaches in optical microscopy. The *Google Colab* notebook that was created as part of this work can be used by microscopist trainees to develop their skills and intuition for STED microscopy before using the microscope for the first time. The optimal set of parameters defined in `pySTED` for a specific fluorophore can guide the parameter choice on a real microscope, but should not replace optimization in real experimental settings to account for environmental effects and biological variability.

The simulation platform was built to be versatile and modular. This allows the users to create and test the efficiency of AI-strategies and adaptive imaging scheme before deploying them on a real microscope. For instance, both DyMIN [8] and RESCue [60] microscopy are readily available to the users. Additionally, the community can contribute open-source modules that would meet their experimental settings.

Smart-microscopy requires the development of tools and modules to increase the capabilities of the microscopes [10, 61] which can be challenging when working on a real microscopy system. The development of simulation software is one way to mitigate the difficulty of building an AI-assisted microscopy setup. We mainly focused on the selection of imaging parameters which is one branch of AI-assisted microscopy but also showed that `pySTED` can be successfully applied to data augmentation in supervised learning settings. A recent trend in microscopy focuses on the implementation of *data-driven* microscopy systems. For example, systems are built to automatically select informative regions or improve the quality of the acquired images [62, 63]. The development and validation of such *data-driven* systems could be achieved with `pySTED`. An interesting avenue to pursue for *data-driven* systems could rely on generative models to create diverse datamaps on-the-fly instead of relying on existing databanks of STED microscopy images, which could be integrated to the modular structure of the `pySTED` simulation environment. Online ML optimization strategies tested in the `pySTED` environment showed similar performances when transferred to the real microscopy environment, opening new possibilities to characterize and benchmark novel *data-driven* microscopy approaches in `pySTED` prior to their deployment on real biological samples.

We also tackle the training of an RL agent, the first for optical microscopy, which would be impossible without the access to a large databank of simulated data. The RL agent enables a full automatization of the imaging parameter selection on a real system when deployed from `gym-STED`, an OpenAI `gym` environment built around `pySTED` [52]. Domain randomization was used heavily within the simulation platform [54] which resulted in a RL agent that could adapt its parameter selection to a wide variety of experimental conditions, even in living samples. Such strategies could be transformative to democratize STED microscopy to a larger diversity of experimental settings and allow non-expert users to acquire high-quality images on a new sample without previous optimization sessions.

While RL agents can represent a powerful tool to automatize microscopy setups, they must be trained on a very large number of examples (e.g. 12M steps in this work) [17, 30], which would be infeasible on a real microscopy setup. The `pySTED` simulation environment allowed the RL agent to bridge the gap between simulation and reality without requiring any fine-tuning. This makes `pySTED` an appealing platform for RL development as it is particularly well suited for complex control tasks requiring temporally distinct trade-offs to be made [20]. In this work, the model relied on a constant preference function to convert the multi-objective optimization into a single reward function. This preference function is ultimately user-dependant. This could be complemented in the future by incorporating RL from human-feedback in the training of the RL model [64, 65]. In future work, temporal dynamics could also be implemented in `pySTED` to open new possibilities to fully automatize the selection of informative regions and of imaging parameters in an evolving environment.

## Acknowledgments

Sarah Pensivy and Tommy Roy from the Neuronal Cell Culture Platform of the CERVO Brain Research Center for the preparation of the dissociated hippocampal cultures. Valérie Clavet-Fournier for immunostainings of neuronal proteins. Mélissa Bilodeau for the help with the `pySTED` logo. Funding was provided by grants from the Natural Sciences and Engineering Research Council of Canada (NSERC) (RGPIN-06704-2019 to F.L.C.), Fonds de Recherche Nature et Technologie (FRQNT) Team Grant (2021-PR-284335 to F.L.C and A.D.), the Canadian Institute for Health Research (CIHR) (F.L.C.), and the Neuronex Initiative (National Science Foundation 2014862, Fonds de recherche du Québec - Santé 295824) (F.L.C.). A.D. is a CIFAR Canada AI Chair and F.L.C. is a Canada Research Chair Tier II. A.B. was supported by a scholarship from NSERC. A.B. and A.M.G. were awarded an excellence scholarship from the FRQNT strategic cluster UNIQUE.

## Author contributions

A.B., A.D. and F.L.C. designed the study. A.B., A.M.G., B.T., A.D. and F.L.C. developed the `pySTED` simulation platform. A.B. performed the DL and RL experiments and evaluated their performance. A.B. and A.M.G. evaluated the performance of the `pySTED` platform and designed the ML experiments. A.B.,

J.C. and J.H. performed the live-cell and two-color experiments. A.B. evaluated the deployment of the ML and RL strategies in real experimental settings. A.B. and F.L.C. wrote the manuscript.

## Data availability

All the datasets used to train the models and the corresponding acquired images are available for download at https://s3.valeria.science/flclab-pysted/index.html.

## Code availability

All code used in this manuscript are open source. Code for the STED simulation platform, `pySTED`, is available from https://github.com/FLClab/pySTED. The bandit algorithms and training procedures are available from https://github.com/FLClab/optim-sted. The `gym-STED` environment and training routines are available from https://github.com/FLClab/gym-sted-pfrl and https://github.com/FLClab/gym-sted respectively.

## 4 Methods

### 4.1 pySTED simulation platform

Two main software implementations are incorporated within the `pySTED` simulation platform: i) point spread functions (PSF) calculation, and ii) emitter-light interactions.

#### PSF calculation

PSF calculation in `pySTED` is inspired by previous theoretical work from Leutenegger *et al*. [1] and Xie *et al*. [2] (Figure 1b). As in Xie *et al*. [2], we calculate the excitation and depletion PSF by using the electric field (Figure 1b). The Effective PSF (E-PSF) is calculated by combining the excitation, depletion and detection PSFs using

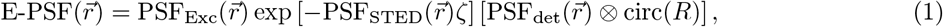

where *R* is the radius of the imaged aperture [3] and *ζ* is the saturation factor of the depletion defined as 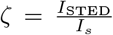 with *I*_*s*_ being the saturation intensity [1]. The left-hand side of equation 1 represents the probability that an emitter at position 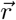 contributes to the signal [4] and is calculated in `pySTED` using *ηp*_exc_ with

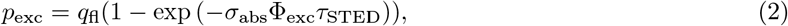

where *q*_fl_ is the quantum yield of the fluorophore, *σ*_abs_ the absorption cross-section, Φ_exc_ the photon flux from the excitation laser and *τ*_STED_ the period of the STED laser. The *η* parameter allows the excitation probability to be modulated with the depletion laser or allows time-gating to be considered during the acquisition [1, 5]. Time-gating consists in activating the detector within a small window of time (*T*_g_, typically 8 ns) after the excitation pulse (*T*_del_, typically 750 ps) to prominently detect photons coming from spontaneous emission. The simulations performed with `pySTED` follow the scheme of pulsed-STED microscopy in which time-gating mostly reduces correlated background [5]. Following the derivation from Leutenegger *et al*. [1] and assuming that *T*_g_ ≥ *τ*_STED_, the emission probability of a fluorophore is described as

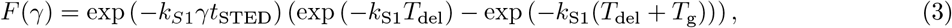

where *k*_S1_ is the spontaneous decay rate,, is the effective saturation factor, 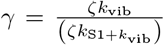 with *k*_vib_ the vibrational relaxation state of *S*_0′_ and *t*_STED_ is the STED pulse width (Figure 1c,d). In the confocal case (*I*_STED_ = 0), the emission probability simply reduces to

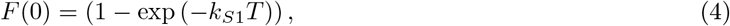

where *T* is the period between each STED pulses. This allows the probability of spontaneous decay *η* to be calculated using *F*(*γ*)/*F*(0). The calculated E-PSF is convolved on the datamap to simulate the photons that are emitted and the one measured by the detector.

In real experiments, the number of detected photons is affected by several factors (e.g. photon detection and collection efficiency of the detector, the detection PSF, the fluorophore brightness, etc.), which were also integrated in the `pySTED` simulation environment (Supplementary Tab 1-5). We also included the possibility to add typical sources of noise that occur in a targeted microscopy experiment such as shot noise, dark noise, and background noise which are all modeled by Poisson processes (Supplementary Tab. 3).

#### Emitter-light interactions

In a real microscopy experiment, the emitters can be degraded as they interact with the excitation or depletion light. Photobleaching is the process by which an emitter becomes inactive following light exposure [6]. In STED microscopy, this process is mainly caused by the combination of the excitation and depletion laser beam [6]. Reducing photobleaching is an optimization objective that the microscopist has to target during an imaging session and that must be minimized to preserve sample health and sufficient imaging contrast. Hence, we implemented a realistic photobleaching model within the `pySTED` simulation software. The photobleaching model is based on the derivations from Oracz *et al*. [6] which were validated on real samples. Figure 1d presents the energy states, the decay rates, and the photobleaching state, *β* that are used within the photobleaching model.

As in Oracz *et al*. [6], we define the photobleaching rate as

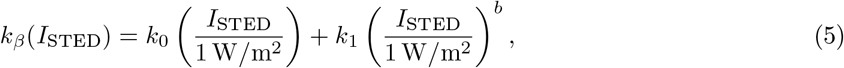

where *k*_0_, *k*_1_ and *b* are dependant on the fluorophore and have to be determined experimentally. In the default parameters of `pySTED` we assume that the linear photobleaching term is null (*k*_0_=0) and that photobleaching occurs only from S1 during the STED pulse. Other photobleaching parameters could be easily integrated considering the modular structure of `pySTED`. We define the effective photobleaching rate *k*_*b*_ as the number of emitters transitioning from the S1 state to the photobleached state (*P*_*β*_) over the course of a laser period *T*

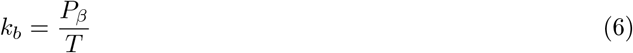

with

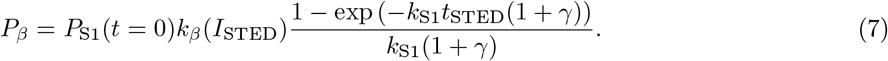

In `pySTED` the number of emitters *N* in a pixel is updated by calculating their survival probability *p* = exp (−*k*_*b*_*t*) from a Binomial distribution for a given dwelltime *t* (Figure 1e). While most parameters can be obtained from the literature for a specific fluorophore, some parameters such as *k*_1_ and *b* need to be determined experimentally [6]. Given some experimental data (or *a priori* about the expected photobleaching of a sample) we can estimate the photobleaching properties (*k*_1_ and *b*) of a fluorophore with

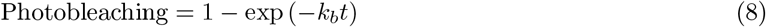

by using non-linear least-squares methods. We can also apply a similar process to estimate the absorption cross-section (*σ*_*abs*_) of a fluorophore to optimize the confocal signal intensity to an expected value Oracz *et al*. [6].

### 4.2 Realistic datamaps

A realistic datamap, that can be used in `pySTED`, is generated by predicting the position of emitters in a real super-resolved image. A U-Net model (U-Net_datamap_, implemented in PyTorch [7]) is trained to predict the underlying structure of a super-resolved image (Supplementary Tab. 18). A single U-Net_datamap_ was trained in this manuscript with images of different sub-cellular structures (F-actin, Tubulin, PSD95, αCaMKII, and LifeAct) and was used to generate all datamaps to train and validate the ML, DL, and RL models presented in this study. U-Net_datamap_ has a depth of 4 with 64 filters in the first double convolution layer. Padding was used for each convolution layer to keep the same image size. As in the seminal implementation of the U-Net [8], maxpool with a kernel and stride of 2 was used. The number of filters in the double convolution layers doubled at each depth. In the encoder part of the model, each convolution is followed by batch normalization and a Rectified Linear Unit (ReLU). Upsampling is performed using transposed convolution. The decoder part of the model uses double convolution layers as in the encoder part of the model. At each depth of the model, features from the encoder are propagated using skipping links and concatenated with the features obtained following the upsampling layer. A last convolution layer is used to obtain a single image followed by a sigmoid layer.

As previously mentioned, the goal of the U-Net is to predict the underlying structure of super-resolved images. Training U-Net_datamap_ in a fully-supervised manner requires a training dataset of associated super-resolved images and underlying structures. However, such a dataset does not currently exist. Mathematically, a microscopy image is obtained from the convolution of the microscope E-PSF with the position of fluorophores at the sample. In the images from Durand *et al*. [9], the E-PSF of the microscope can be approximated by a Gaussian function with a full width at half maximum of ∼ 50 nm. Hence, U-Net_datamap_ can be trained to predict the datamap that once convolved with the E-PSF will be similar to the input image (Figure 1f). The *L*_2_ error is calculated between the Gaussian convolved datamap and the original input image as the loss function to minimize.

To train the model we used good quality STED images of diverse neuronal proteins from an existing dataset [9] (quality > 0.7). In Durand *et al*. [9], the quality score of an image was obtained by asking an expert to rate the image based on a qualitative assessment of the resolution of the structure of interests and the signal to noise ratio on a scale from 0 to 1. The quality scores from the original dataset were used to train a deep learning model to automatically rate the quality of an image. Supplementary Tab. 7 presents the proteins imaged and the number of images that were used for training. Each 224×224 pixels image is augmented with three (3) 90° rotations. The Adam optimizer was used with default parameters using a learning rate of 1× 10^−4^. The model was trained for 1000 epochs with a batch size of 32. We selected the model with the best generalization properties on the validation set, obtained from the mean squared error between the input image and the datamap after applying the Gaussian convolution.

By default, the predicted datamap reconstructs the background noise from the image. Filtering can be applied on the predicted datamap to reduce the impact of noise. The number of emitters can be adapted to the experimental context which is then converted into an integer value. U-Net_datamap_ was trained with 224 × 224 pixels images but images of arbitrary size can be processed at inference time.

#### Datamaps from deconvolution

Datamaps were generated using the Richardson-Lucy deconvolution implementation from van der Walt *et al*. [10]. The E-PSF of the input image were approximated by a Gaussian function with a full width at half maximum of ∼50 nm. 30 iterations were used for the deconvolution algorithm.

### 4.3 Imaging optimization objectives

#### Resolution

We calculated the resolution of the images by using the parameter-free image resolution estimation based on decorrelation analysis that was developed by Descloux *et al*. [11]. Decorrelation analysis was used due to its simplicity in transferring from simulation to real imaging conditions.

#### Photobleaching

In all experiments involving the photobleaching as one of the imaging optimization objective, we measured the loss of the fluorescence signal between a low-resolution image that is acquired before (Confocal 1) and after (Confocal 2) the high-resolution (STED) acquisition [9]. The photobleaching is defined as

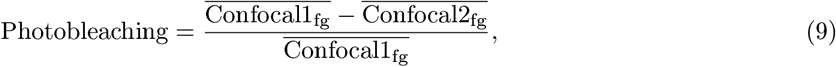

where 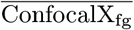 is the average signal on the foreground of the first confocal image (Confocal1). The foreground mask is determined using an Otsu threshold on the Confocal1.

#### Signal Ratio

We calculate the signal ratio as the ratio between the intensity in the high-resolution image and the respective confocal image using the following equation

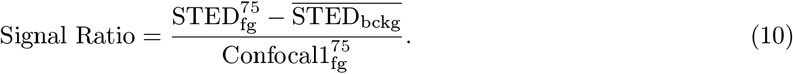

The foreground mask of the STED and confocal images are determined using the Otsu method. The fore-ground signal in the mask is calculated as the 75^th^ percentile of the image (STED or Confocal1). 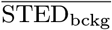 represents the mean signal of the background signal in the STED image.

#### Artefact

We measured imaging artefacts with a metric inspired by SQUIRREL [12] and MASK-SSIM approaches [13]. Specifically, we map the super-resolution image (SR) to a low-resolution image using a similar procedure to SQUIRREL but compare structures only within a foreground mask. This foreground mask is obtained using the Otsu method. The average structural similarity index (SSIM) on the foreground between the low-resolution and the optimized SR image is reported as the metric. The value of the artefact metric that is reported in the paper is

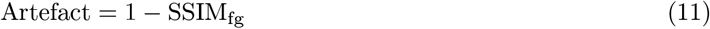

Artefact =1 - SSIM_fg_

### 4.4 Comparison of pySTED simulations with real acquisitions

We compared `pySTED` simulated images with images acquired on a real STED microscope with similar imaging parameters. To evaluate the reliability of the simulations, we acquired 10 images using a different combination of parameters. We varied each imaging parameter over a range that is commonly used for routine STED experiments and that would not damage the microscope (e.g. detectors, see parameters in Supplementary Tab. 6). We used a sample of immunostained cultured hippocampal neurons of the neuronal protein Bassoon tagged with the fluorophore ATTO-647N. The small clusters formed by Bassoon are well suited for measurements of resolution. The same parameter combination is used in `pySTED` and on the microscope for fair comparison.

We optimized the photobleaching constants (*k*_1_ and *b*) and the STED cross-section (*σ*-_STED_) of the fluorophore to match the measured photobleaching and resolution values using a least-squares method (data from Supplementary Fig. 4a, right). It is implemented iteratively, with the optimization of photobleaching and resolution done sequentially and repeated 15 times, since the optimization of *σ*-_STED_ also affects photobleaching.

For each acquired real STED images, a datamap is predicted with the U-Net_datamap_. The number of emitters per pixel is obtained by multiplying the datamap with a correction factor *f* to match the fluorescence signal in the real images. This correction factor *f* can be obtained by fitting the intensity value obtained at pixel (*x, y*) to the real intensity of the acquired confocal image (*I*_CONF_(*x, y*)). The intensity value of the synthetic image is approximated as the product between the E-PSF and the datamap (*D*)

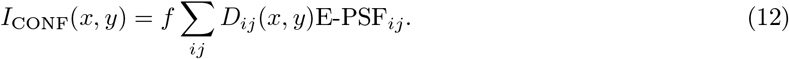

### 4.5 Weakly supervised learning for the segmentation of F-actin nanostructures

We compared three training schemes (5 random initializations per scheme) to train a U-Net model to segment two F-actin nanostructures (Fibers and Rings): i) original model from Lavoie-Cardinal *et al*. [14] (O), ii) a model that uses a quantile normalization of the large image (min/max normalization using the 1^st^ and 99^th^ quantile) and increased data augmentation during training (N, see below), iii) and a model trained as in ii) with synthetic images (N+S). In all conditions, the same architecture, training procedure, and dataset are used following the methods from Lavoie-Cardinal *et al*. [14]. A model is trained for 700 epochs and the best model on the validation dataset is kept for testing. To compare only the impact of the training, the validation dataset is kept constant in all training instances.

In all training schemes, an augmentation has a 50% probability of being selected. For the training scheme O [14], the augmentations consisted of horizontal/vertical flips, intensity scale, and gamma adaptation. For the approaches using an increased data augmentation scheme (N and N+S), the augmentations from O are combined with random 90° rotations, crop normalization (1^st^ and 99^th^ percentile) and more intensity scale and gamma adaptation operations.

#### Synthetic F-actin images

The U-Net_datamap_ model (Figure 2) was used to extract the datamaps of all valid crops in the training dataset (contains >10% of dendrite, 256× 256 pixels, 25% overlap). Five synthetic images with different resolution and noise properties were simulated for each crops with `pySTED` using a parameters combination that would minimally allow to resolve the F-actin nanostructures (Figure 2b and Supplementary Tab. 8).

#### Generation of subsets

Models (with constant parameter initialization) were trained on subsets of the original dataset, to evaluate if `pySTED` can help reduce the number of original images in the training dataset. Five subsets with 0.75, 0.5, 0.25, 0.1, 0.05, 0.025 ratios were used for training. A ratio of 0.025 corresponds to training on a single image (42 images in training dataset). When an image is discarded from a subset, its corresponding crops (synthetic included) are also removed from training (Supplementary Fig. 7).

#### Performance evaluation

The average precision (AP) is used for performance evaluation. The AP is obtained from the precision and recall measured by the predicted segmentation compared to the ground truth manual annotations. The AP corresponds to the area under the *p*_*∗*_(*r*) curve. *p*_*∗*_(*r*) is given by the maximum precision value that can be attained at any recall *r*_*i*_ greater than recall *r, i*.*e*.

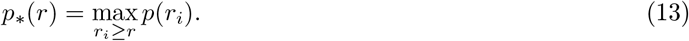

The AP is calculated as

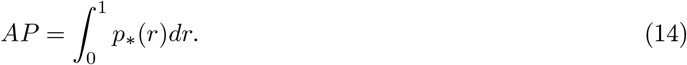

### 4.6 Multi-objective bandit optimization

The multi-objective bandit optimization aims at finding a set of imaging parameters that simultaneously optimizes all imaging optimization objectives. Such a multi-objective problem is ill-defined as there exists a set of Pareto optimal objectives that could be used to solve the task. Hence, an external input, *e*.*g*. a microscopist, is required to make the necessary trade-offs over the course of the optimization session.

#### 4.6.1 Algorithms

The goal of the algorithm is to learn the mapping between the imaging parameters (*e*.*g*. laser power, pixel dwelltime) and the imaging optimization objective (*e*.*g*. resolution, photobleaching, artefact or signal ratio) by exploring the parameter space while exploiting its current knowledge of the parameter space to acquire high-quality images. A single model is built for each optimization objective as in Durand *et al*. [9]. The exploration/exploitation trade-off is achieved via Thompson Sampling (TS) [15]. At each time step of the optimization, a function is sampled from the posterior of each model. The expected imaging optimization objective associated with each imaging parameters options are combined. The preferred combination is selected and an image is acquired with the associated parameters. The imaging optimization objectives are calculated from the resulting image and used to update each model.

The range of imaging parameters was normalized in [−1, 1] using a min-max normalization. The min-max values were given from the pre-defined range of a parameter. All models are trained from scratch. At the start of each optimization session, 3 images are acquired with parameter combinations obtained from expert knowledge allowing the models to gain insights about the imaging task. For further implementations, these parameter combinations could be obtained from i) a previous imaging session, ii) different fluorophore, or iii) publications from the field.

##### Kernel-T

Kernel-TS is implemented by following the procedure from Durand *et al*. [9]. The regression model that maps the imaging parameters to the imaging optimization objectives is a non-parametric Gaussian Process. All of the parameters of the method (*e*.*g*. the kernel bandwidth or bounds on noise) were based on the recommendations from the original manuscript [9]. Kernel-TS works on a discrete parameter space of 10 points for each optimized parameters. The values of imaging optimization objectives are rescaled using a whitening transformation.

##### LinTSDiag

LinTSDiag is a neural network implementation of TS [16]. LinTSDiag was previously implemented to solve a 2 parameter DyMIN task [17] (Supplementary Tab. 19). In this implementation, the neural network is a fully connected network with 2 layers of hidden sizes of 32. After each layer ReLU activation is used and followed by a dropout layer (probability of 0.2). The last layer of the model projects to a single imaging optimization objective value. The model is implemented in PyTorch [7] and relies on the seminal implementation from Zhang *et al*. [16]. The loss of the model is the mean squared error and is optimized using stochastic gradient descent with a learning rate of 1×10^−3^. After each acquisition, the weights of the model are updated until the error is < 1× 10^−3^ or 1000 updates have been done. During training, the imaging optimization objectives are rescaled into a [0, 1] range.

Two parameters (*ν* and *λ*, see Zhang *et al*. [16]) control the exploration of the model. Increasing their values results in more exploration. In all experiments using LinTSDiag A = 0.1 is used. The parameter *ν* varied depending on the task: i) in simulation *ν* = 0.01 (Figure 3a-c), ii) in 4 parameter optimization *ν* = 0.1 (Figure 3e-f), and iii) in 6 parameter optimization *ν* = 0.25 (Figure 3g-h).

LinTSDiag handles continuous parameter space. Hence, it is not possible to display all of the possible trade-offs. To reduce the number of possibilities, only the Pareto optimal combination of optimization objectives are displayed (Pareto front). The Pareto optimal options are extracted using NSGA-II [18] with the implementation from the DEAP python library [19]. Since computing the Pareto front is computationally expensive, a stopping criterion is used to reduce the calculations as previously reported by Roudenko & Schoenauer [20]. The stopping criterion is based on the rolling standard deviation of the maximum crowding distance (window size of 10) during the NSGA-II search. The search is stopped when the standard deviation is lower than 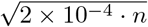, where *n* is the number optimization objectives [20]. In theory, the NSGA-II search should restart from scratch after each acquisition. However, given the high dimensionality of the parameter space, this may lead to high variability in the proposed parameter combination. To reduce this variability, Deb *et al*. [21] proposed to keep a fraction of the previous options as a warm start of the NSGA-II search. In this work, 30% of the previous options are randomly sampled and used as starting points for the next NSGA-II search. The resulting Pareto front of imaging optimization objectives is shown to the preference articulation method.

##### Contextual LinTSDiag

The contextual version of LinTSDiag heavily relies on the implementation of LinTSDiag described above (Supplementary Tab. 20). In this work, the contextual information was used to solve a DyMIN microscopy task. As previously mentioned the confocal image serves as contextual information, but any other contextual information pertinent to the task could be provided to the model. The confocal image is encoded with a 2 layers convolutional neural network. A first convolution layer with 8 filters, kernel size of 3 and padding of 1 is followed by a batch normalization layer, maxpooling layer (size 4; stride 4) and ReLU activation. A second convolution layer of 16 filters is followed by a batch normalization layer. Global average pooling is used to generate a vector embedding. This is followed by a ReLU activation and dropout layer with a probability of 0.2. The embedding is projected to 32 features using a fully connected layer and is followed by a ReLU activation and a dropout layer with a probability of 0.2. The contextual features are concatenated with the parameter features (described in LinTSDiag). A single-layer fully-connected model with a hidden size of 64 is used to predict the imaging optimization objectives. ReLU activation is used at the hidden layer. A single contextual encoder is created and shared between the imaging optimization objectives. The same training procedure and NSGA-II search are used as in LinTSDiag.

The exploration parameters *λ* = 0.1 and *ν* = 0.25 were used in simulation (Figure 4c-g) and in 3D DyMIN optimization (Figure 4h-j).

#### 4.6.2 Preference articulation

The optimization algorithms output possible trade-offs between the imaging optimization objectives. The preference articulation step consists in selecting the trade-off that is the most relevant for the task. Two preference articulation methods were used in the bandit optimization: manual selection and automatic selection [9].

##### Manual selection

This method requests a manual input from the microscopist at each image acquisition. The microscopist is asked to select the trade-off that is inline with their own preferences from the available options (point cloud). This method was used in all experiments on the real microscope using the bandit optimization scheme (Figure 3e-h and Figure 4h-j).

##### Automatic selection

This method aims at reducing the number of interventions from the microscopist in the optimization loop by learning their preferences prior to the optimization session. In Durand *et al*. [9], the neural network implementation PrefNet was used to learn the preferences from an Expert. In the current work, two PrefNet models were trained from the preferences of an Expert. The same model architecture and training procedure were used as in Durand *et al*. [9]. A first model is trained for the STED optimization to select from the resolution, photobleaching, and signal ratio imaging optimization objectives. A second model is trained for the DyMIN optimization to select the trade-off between resolution, photobleaching, and artefact. The PrefNet model is used to repeatedly make the trade-offs in multiple optimizations in the simulation environment (Figure 3b-d and Figure 4c-g).

### 4.7 Reinforcement learning experiments

A RL agent interacts with an environment by sequentially making decisions based on its observations. The goal of the agent is to maximize its reward signal over the course of an episode.

#### RL formulation

The general problem in RL is formalized by a discrete-time stochastic control process, *i*.*e*. it satisfies a Markov Decision Process (MDP). An agent starts in a given state *s*_*t*_ ∈ 𝒮 and gathers some partial observations *o*_*t*_∈ 𝒪. In an MDP, the state is fully observable, that is the agent has access to a complete observation of a state *s*_*t*_. At each time step *t*, the agent performs an action *a*_*t*_ ∈𝒜 given some internal policy π after which the agent receives a reward *r*_*t*_ ∈ ℛ and transitions to a state *s*_*t*+1_ ∈ 𝒮 with a state transition function 𝒯 (*s*_*t*+1_ |*s*_*t*_, *a*_*t*_). Following the state transition, a reward signal *r*_*t*_ = *R*(*s*_*t*_, *a*_*t*_, *s*_*t*+1_) is provided to the agent as feedback. The goal of the agent is to maximize the cumulative reward over the trajectory *τ* = (*s*_*t*_, *a*_*t*_, *s*_*t*+1_, *a*_*t*+1_, …). Formally, the cumulative reward may be written in the form of

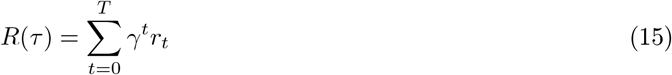

where *γ*, is a discount factor in the range [0, 1] to temporally weight the reward. Intuitively, using a discount factor close to 1 implies that the credit assignment of the current action is important for future reward, which is the case for long planning horizon, while a discount factor close to 0 reduces the impact of temporally distant rewards [22].

#### Reward function

The optimization of super-resolution STED microscopy is a multi-objective problem (e.g. Resolution, Signal Ratio, and Photobleaching). However, the conventional RL settings and algorithms assume the access to a reward function that is single-valued, in other words a single-objective optimization [22]. Several methods were introduced to solve the multi-objective RL setting, for instance by simultaneously learning multiple policies or by using a scalarisation function (see Hayes *et al*. [23] for a comprehensive review). The scalarisation function is simple to implement and allows all of the algorithms that were developed for RL to be used, but assumes that the preference from the user are known a priori. In this work, the multi-objective RL setting was transformed into a single scalar reward by using the neural network model, PrefNet [9], that was developed in the bandit experiments. Indeed, the PrefNet model was trained to reproduce the trade-off that an expert is willing to make into the imaging optimization objective space. The PrefNet model does so by assigning a value to a combination of imaging optimization objectives. The values predicted by the model for a combination of optimization objectives are arbitrary but the ranking of these values is accurate. Hence, the values from the PrefNet model is proportional to the image quality. The reward of the agent can then be defined using equation 16. For safety precautions when deploying the agent on a real microscopy system, the agent incurs a reward of -10 when the frequency of photons on the detector is higher than 20 MHz.

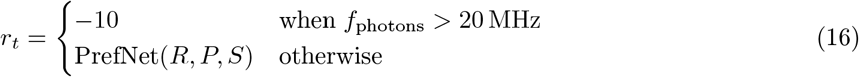

While the negative reward can be used to limit the selection of actions that could damage the microscope, it is not required. For instance, the results showed in Figure 6d-e and Extended Fig. 4 used a version of the reward function that did not include the negative reward. It is worth noting that in these cases, the range of parameters should be carefully selected to avoid damages to the microscope.

#### Agent

The Proximal Policy Optimization (PPO) model [24] was used for all RL experiments. PPO is considered state-of-the-art for many control tasks, and is widely used in robotics [25]. PPO allows continuous action space making it suitable for the task of microscopy parameter tuning. It is an on-policy algorithm meaning that the same policy is used during the data collection and the updating phases. The model uses a deep neural network to map the state to the actions. Since PPO is an actor-critic method, it simultaneously learns a policy function and a value function that measures the quality of a selected action (Supplementary Tab. 21 and 22). Both functions use the same model architecture. A convolutionnal neural network (CNN) extracts information from the visual inputs and a linear neural network (LN) extracts information from the history of the episode. The CNN encoder is similar to the one used in Mnih *et al*. [26]. The encoder is composed of 3 layers of convolutions each followed by a leaky ReLU activation. The kernel size of each layer is 8, 4, 3 with a stride of 4, 2, 1. This allows the spatial size of the state space to be reduced. The LN model contains 2 linear layers projecting to sizes 16, 4. The information from both layers is concatenated and mapped to the action space using a LN layer.

During training, the Adam optimizer is used with default parameters and a learning rate of 1×10^−4^. The batch size of the model is set at 64. Each 512 steps in the environment, the model is trained for 10 batches which are randomly sampled from the previous 512 steps. A maximal gradient of 1.0 during backpropagation is used to stabilize training.

#### Synthetic Datamaps

A bank of datamaps was generated using U-Net_datamap_. Supplementary Tab. 11 presents the number of images per structures that were available during training. Datamaps were randomly cropped to 96 × 96 pixels with a higher probability of being sampled within the foreground of the datamap.

Random data augmentation is performed online with a 50% probability: random [1, 3] 90° rotations, up-down flips, and left-right flips. The resultant cropped datamap is multiplied by a value that is sampled from *𝒩* (*µ* = 40, *a*- = 4) and turned into an integer array using the floor operation.

#### Synthetic Fluorophores

Synthetic fluorophore properties are generated on-the-fly during training by uniform sampling. Supplementary Tab. 12 displays the range of possible fluorophore properties. The parameters *k*_1_, *b*, and *σ*_abs_ are optimized using the procedure described in section 4.4. During the optimization it is assumed that the maximal number of emitters is 40. A scaling factor that is dependant on the type of structure is used during the optimization (Supplementary Tab. 13). At each iteration, the fluorophore parameters from the previous iterations are used as a starting point. The initial conditions of parameters are *k*_1_ = 2.9 × 10^−16^, b = 1.66, and *σ*_abs_ = 3.2 × 10^−21^ m^2^.

### 4.8 STED microscopy experiments

#### 4.8.1 STED-imaging

Super-resolution imaging of neuronal proteins was performed on an Abberior Expert-Line STED system (Abberior Instruments GmbH, Germany) equipped with a 100x 1.4NA, oil objective lens (Olympus, UPLSAPO100XO), motorized stage and auto-focus unit. Far-red dyes were imaged using a 640 nm pulsed diode (40 MHz), a 775 nm depletion laser (40 MHz) and a ET685/70 (Chroma, USA) fluorescence filter. Fluorescence was detected on an avalanche photodiode detectors (APD) with approximately 1 Airy unit detection pinhole. Images were processed using FIJI (ImageJ) software. Single- and two-channel imaging of tubulin, NPC, and Golgi in Vero cells was performed on an Infinity line microscope (Abberior Instruments GmbH, Germany) using imaging settings as described in Heine *et al*. [27].

Prior to the optimization, the excitation power of the confocal acquisition needed to be set to acquire < 200 photons in 10 µs. To do so, the excitation power was first set to 10 µW and was halved until this criterion was met. This value is used by the model to incorporate knowledge about the brightness of the sample.

##### Kidney epithelial cell culture

Vero B4 cells were obtained from the DSMZ-German Collection of Microorganisms and Cell Cultures were maintained in DMEM (Gibco) supplemented with GlutaMAX (Thermo Fisher Scientific), 10% FBS (Sigma-Aldrich), 1 mM sodium pyruvate (Thermo Fisher Scientific), and Penicillin-Streptomycin (100 µl/ml and 0.1 mg/ml; Sigma-Aldrich) at 37°C with 5% CO2.

##### Sample preparation and staining procedures

For indirect immunostaining, cells were fixed in 8% paraformaldehyde (PFA) in phosphate-buffered saline (PBS) and permeabilized with 0.5% Triton X-100/PBS for Nuclear Pore Complex (NPC) proteins (Mab414, 1:200; abcam, code: ab24609) and Golgi (Giantin, 1:200; abcam, code: ab80864) staining. Methanol was used as a fixative for Tubulin staining (1:500; abcam, code: ab18251).

After blocking with 2% BSA/0.1% Tween20/PBS, cells were incubated with the primary antibody for 1 hour at the specified dilutions. Detection of primary antibodies was achieved using secondary STAR RED goat anti-mouse IgG (1:200, abberior GmbH, code: STRED-1001-500UG) and STAR ORANGE goat anti-rabbit IgG (1:200, abberior GmbH, code: STORANGE-1002-500UG) for double staining of the NPC and Golgi. Tubulin was labeled with STAR RED goat anti-rabbit IgG (1:200, abberior GmbH, code: STRED-1002-500UG). Secondary antibodies were also incubated for 1 hour.

After stringent washing with PBS, cells were mounted in MOUNT SOLID ANTIFADE (abberior GmbH, code: MM-2013-2X15ML). Protocol was adapted from Wurm *et al*. [28].

#### 4.8.3 Neuronal cell culture

Neuronal cultures from the hippocampus were obtained using neonatal Sprague Dawley rats, adhering to the animal care guidelines set by Université Laval. The rats, aged P0-P1, were sacrificed through decapitation before the hippocampi were dissected. The cells were then seeded onto 12 and 18 mm coverslips coated with poly-d-lysine and laminin, for fixed (12 mm, 40,000/coverslip) and live-cell (18 mm, 100,000 cells/coverslip) STED imaging. Neurons were cultivated in a growth medium composed of Neurobasal and B27 (in a 50:1 ratio), enriched with penicillin/streptomycin (25 U/mL; 25 µg/mL) and 0.5 mM L-GlutaMAX (by Invitrogen). Ara-C (5 µM; from Sigma-Aldrich) was added into the medium after five days to limit the proliferation of non-neuronal cells. Twice a week, ∼50% of the growth medium was replaced with serum- and Ara-C–free medium. Cells were used between *Days In Vitro* (DIV) 12-16 for experiments.

##### Sample preparation and staining procedures

Fixation was performed for 10 minutes in 4% PFA solution (PFA 4%, Sucrose 4%, Phosphate Buffer 100mM, Na-EGTA 2mM). Neurons were permeabilized with 0.1% Triton X-100 and aspecific binding sites were blocked for 30 min with 2% goat serum in PBS 20 mM. Primary and secondary antibodies were successively incubated for 2h and 1h respectively. Phalloidin was incubated for 1h. All incubations were done at room temperature, in the blocking solution. Immunostained coverslips were mounted in Mowiol-DABCO for imaging. F-actin was stained with phalloidin-STAR635 (Sigma Aldrich, cat. 30972-20µg, 1:50 dilution). All antibodies used in this study with associated concentrations are provided in Supplementary Tab. 16 & 17.

For the live experiment (Figure 6d), the neurons were incubated for 8 min with SiR-actin (0.8 µM, Cytoskeleton, cat. CY-SC001) diluted in HEPES buffered artificial cerebrospinal fluid (aCSF, in mM: NaCl 98, KCl 5, HEPES 10, glucose 10, CaCl_2_ 0.6, MgCl_2_ 5). For live-cell STED microscopy, coverslips were mounted on a QR chamber (Wagner Instruments, cat. 61-1944) and imaged in HEPES buffered aCSF 5 mM Mg^2+^ / 0.6 mM Ca^2+^ using a gravity perfusion system.

#### 4.8.4 Quantification of biological structures

##### F-actin

Line profiles of ∼1 µm were manually extracted from each image. A linewidth of 3 pixels was used to average the profile values. The autocorrelation function (`statsmodels` library [29]) was calculated from the intensity profile. The length of periodicity of the signal was determined from the first peak maxima.

##### CaMKII-,*β* and PSD95

We segmented clusters using a wavelet segmentation using the implementation from Wiesner *et al*. [30]. The scales used were (1, 2) for STED and (3, 4) for confocal segmentation. A threshold of 200 was used. Small segmentation objects (<3 pixels) were removed and small holes (<6 pixels) were filled. In the STED image segmentation, only the objects part of the confocal foreground were considered. For STED segmentation, watershed was used to split merged segmented objects. The local peak maximum were used as initial seeds. Small segmentation objects (<3 pixels) resulting from the watershed split were filtered out. The properties of each segmented object was extracted using `regionprops` from the `scikit-image` python library [10].

##### TOM20

A similar approach to the one in Wurm *et al*. [31] was used. Briefly, the confocal foreground of each mitochondrion was extracted using the same wavelet segmentation procedure as for CaMKII-,*β* and PSD95. The 2D autocorrelation on square crops of 320 nm x 320 nm centered on each mitochondrion were calculated. The diameter of TOM20 cluster is defined as the standard deviation obtained from a 2D Gaussian curve fit of the autocorrelation profile.

## Extended Figures

**Extended Fig. 1:**
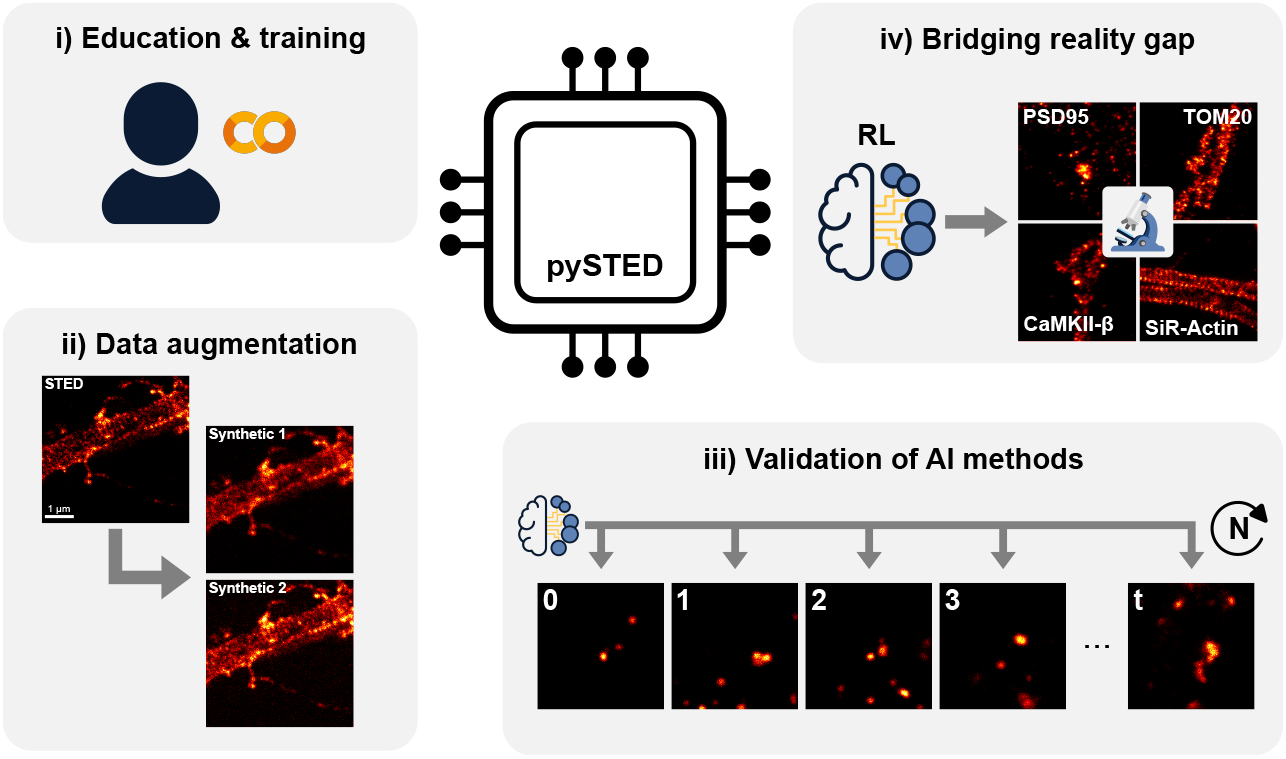
`pySTED` can benefit common microscopy tasks, *i*.*e*. image analysis and acquisition. i) A Google Colab notebook implementing `pySTED` is created for trainees to develop their knowledge and intuition about STED microscopy. ii) `pySTED` can be leveraged in deep learning-related microscopy tasks to artificially augment the training datasets. iii) `pySTED` can be used to develop and thoroughly validate AI methods by limiting the impact of biological variability on the measurements and reducing the biological footprint. iv) `pySTED` reduces the reality gap between simulation and reality by training RL models that learn through interactions with the system. The trained models are then deployed in a wide range of real experimental conditions.

**Extended Fig. 2:**
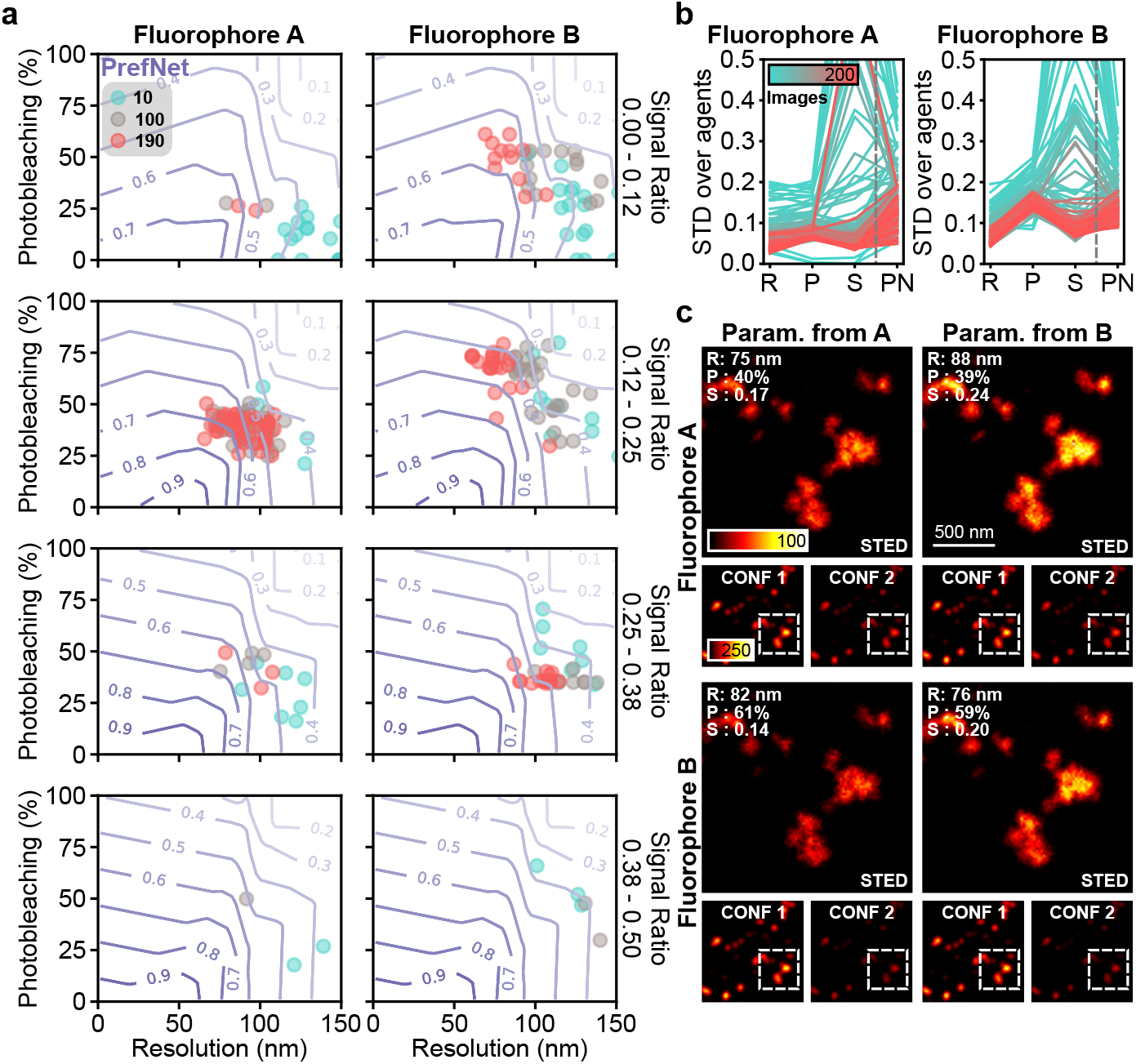
a) Resulting imaging optimization objectives from Kernel-TS at 3 different timesteps (10 - cyan, 100 - grey, and 190 - red) for 50 independent models are presented for increasing signal ratio (top to bottom). With time, Kernel-TS acquires images that have a higher preference score for both fluorophores (purple contour lines) and converges into a similar imaging optimization objective space (red points). b) The standard deviation (STD) of the imaging optimization objectives and of the preference scores decreases during the optimization (cyan to red) supporting the convergence of Kernel-TS in a specific region of the imaging optimization objective space for both fluorophores. The dashed line separates the imaging optimization objectives (R: Resolution, P: Photobleaching, and S: Signal ratio) from the preference network (PN). c) Typical `pySTED` simulations on two different fluorophores (top/bottom) using the optimized parameters on fluorophore A (top) or B (bottom). See Supplementary Tab. 9 for imaging parameters.

**Extended Fig. 3:**
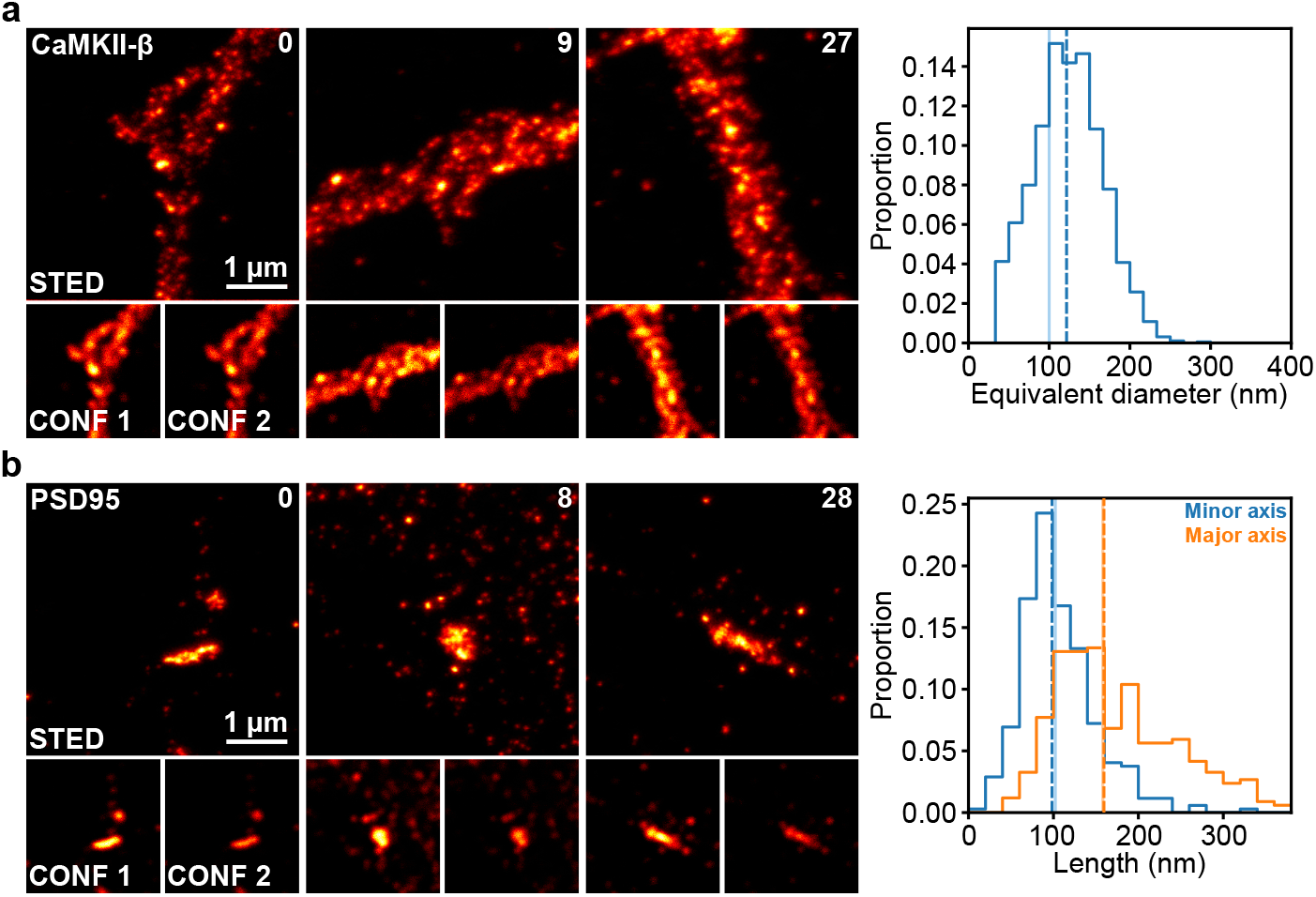
The agent was deployed on a real STED microscope for the imaging of diverse neuronal proteins in dissociated hippocampal cultures. **In distribution** a) (left) The clusters of CaMKII-*β* are revealed in all acquired images. (right) The size of the clusters are extracted from the acquired images (dashed vertical line represents the median of the distribution) and compared to the values that were previously reported in Ferreira *et al*. [1] (solid vertical line). **Out of distribution** b) A bright fluorophore of PSD95 is simulated experimentally (Methods). The STED images reveal the presence of nano-clusters. The minor (blue) and major (orange) axis length of the nano-clusters are measured and compared with the values reported in Nair *et al*. [2].

**Extended Fig. 4:**
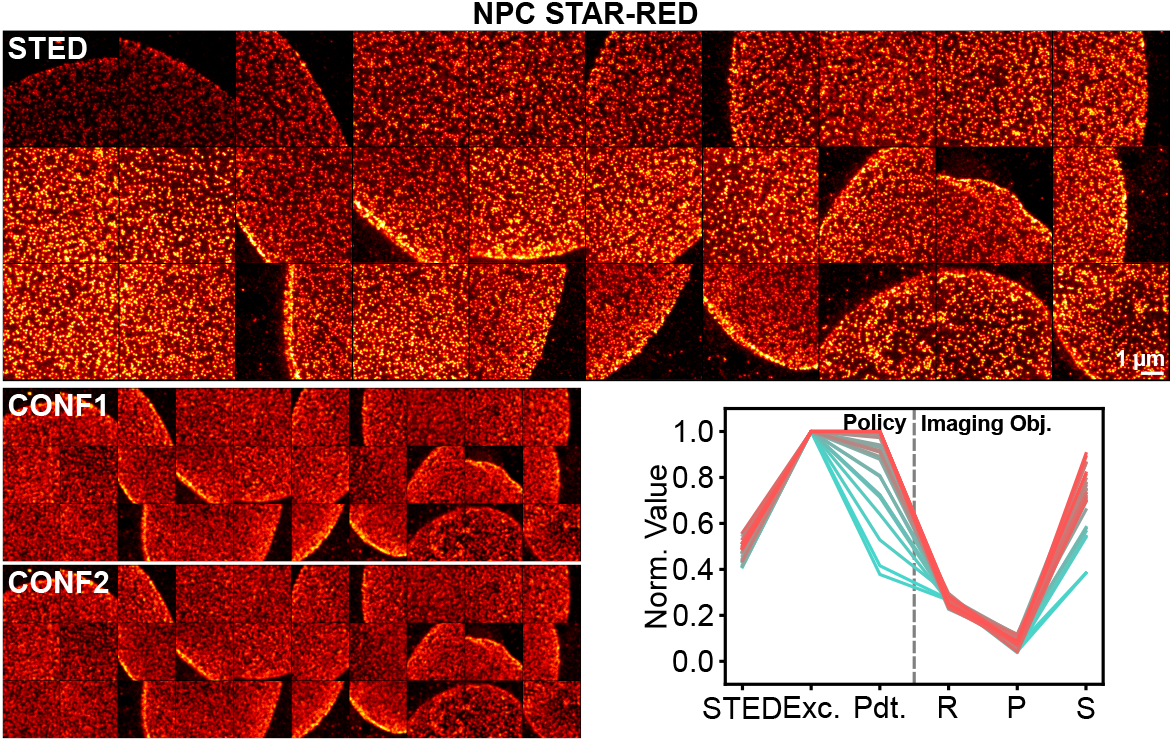
Images acquired by the RL agent in a real experiment on a different microscope. NPC was stained with the STAR-RED fluorophore. The sequence of acquired images goes from top left to bottom right. The confocal images before (CONF1) and after (CONF2) are presented for photobleaching comparison. The CONF1 image is normalized to the CONF2 image. The STED images are normalized to the 99^th^ percentile of the intensity of the CONF1 image. Images are 5.12 µm *×* 5.12 µm. The evolution of the parameter selection (left; STED: STED power, Exc.: Excitation power, Pdt.: Pixel dwelltime) and imaging optimization objectives (right; R: Resolution, P: Photobleaching, S: Signal ratio) are presented.

## Footnotes

1 https://github.com/FLClab/pySTED

