## Supplementary Materials for "Development of AI-assisted microscopy frameworks through realistic simulation in pySTED"

#### Supplementary Material

##### Tables

| Parameter | Description | Default |
|---|---|---|
| $\lambda$ | Wavelength (m) | - |
| Polarization | Phase difference between $x$ and $y$ oscillations (rad) | $\frac{\pi}{2}$ |
| $\beta$ | Beam incident angle (rad) | $\frac{\pi}{4}$ |

Supplementary Tab. 1: Parameters of a `Gaussian Beam`.

| Parameter | Description | Default |
|---|---|---|
| $\lambda$ | Wavelength (m) | - |
| Polarization | Phase difference between $x$ and $y$ oscillations (rad) | $\frac{\pi}{2}$ |
| $\beta$ | Beam incident angle (rad) | $\frac{\pi}{4}$ |
| $\tau_{\text{STED}}$ | Pulse length (s) | $400 \times 10^{-12}$ |
| $\tau_{\text{rep}}$ | Pulse rate (Hz) | $40 \times 10^{6}$ |
| $Z_0$ | Ratio between minimum and maximum intensity ratio (-) | 0 |
| Anti-Stokes | Anti-stokes excitation (bool) | True |

Supplementary Tab. 2: Parameters of a `STED Beam`.

| Parameter | Description | Default |
|---|---|---|
| $n_{\text{airy}}$ | Number of airy disks (-) | 0.7 |
| Noise | Add Poisson noise to signal (bool) | False |
| Background | Average number of photon counts from background (Hz) | 0 |
| Dark count | Average number of photon counts from dark counts (Hz) | 0 |
| PCEF | Photon collection efficiency factor (-) | 0.1 |
| PDEF | Photon detection efficiency factor (-) | 0.5 |
| $T_{\text{del}}$ | Gating delay (s) | $750 \times 10^{-12}$ |
| $T_g$ | Gating time | $8 \times 10^{-9}$ s |

Supplementary Tab. 3: Parameters of a `Detector`. The detector parameters are based on an avalanche photodiode (APD).

| Parameter | Description | Default |
|---|---|---|
| $f$ | Focal length (m) | $2 \times 10^{-3}$ |
| $n$ | Refractive index of the medium (-) | 1.5 |
| NA | Numerical aperture | 1.4 |
| T | Transmission factor (-) | 0.85 |

Supplementary Tab. 4: Parameters of an `Objective Lens`.

| Parameter | Description | Default |
|---|---|---|
| $\sigma_{\text{STED}}$ | Stimulated emission cross-section (m$^2$) | $\sigma_{\text{STED},750} = 4.8 \times 10^{-22}$ [1] |
| $\sigma_{\text{abs}}$ | Absorption cross-section (m$^2$) | $\sigma_{\text{abs},635} = 1 \times 10^{-20}$ $\sigma_{\text{abs},750} = 3.5 \times 10^{-25}$ [1] |
| $\tau$ | Fluorescence lifetime (s) | $3.5 \times 10^{-9}$ [1] |
| $\tau_{\text{vib}}$ | Vibrational relaxation (s) | $1.0 \times 10^{-12}$ [1] |
| $\tau_{\text{tri}}$ | Triplet state lifetime (s) | $25 \times 10^{-6}$ [2] |
| $q_{\text{fl}}$ | Quantum yield (-) | 0.65 [2] |
| $k_0$ | First order term of photobleaching rate (-) | 0 [1] |
| $k_1$ | $b^{\text{th}}$ order term of photobleaching rate (-) | $1.3 \times 10^{-15}$ [1] |
| $b$ | Intersystem crossing rate (s$^{-1}$) | 1.45 [1] |

Supplementary Tab. 5: Parameters of the `Fluorescence`.

| Parameter | Value |
|---|---|
| Excitation power | 3.76 µW |
| Pixel dwelltime | 15 µs |
| Depletion power | 150 mW |
| Gating delay | 750 ps |
| Gating time | 8 ns |
| Line repetitions | 8 |

Supplementary Tab. 6: Default parameters used for the validation of `pySTED` with a real microscope (Supplementary Fig. 3-5).

| Dataset | Images / Total | Training / Validation |
|---|---|---|
| Actin | 1224 / 2669 | 963 (79 %) / 261 (21 %) |
| Tubulin | 309 / 525 (59 %) | 249 (81 %) / 60 (19 %) |
| PSD95 | 307 / 494 (62 %) | 247 (80 %) / 61 (20 %) |
| $\alpha$CaMKII | 280 / 757 (37 %) | 224 (80 %) / 56 (20 %) |
| LifeAct | 246 / 577 (43 %) | 189 (77 %) / 57 (23 %) |
| Total | 2366 / 5022 (47 %) | 1872 (79 %) / 495 (21 %) |

Supplementary Tab. 7: Datasets used to train the U-Net$_{\text{datamap}}$ model to predict the underlying structure [3]. Images with a quality rating $> 0.7$ that were manually obtained from an expert in Durand *et al.* [3] were kept for training.

| Parameter | Range |
|---|---|
| Excitation power | [1, 6] µW |
| Depletion power | [75, 200] mW |
| Pixel dwelltime | [10, 50] µs |
| Background | [0.5, 10]/(50 µs) |

Supplementary Tab. 8: Range of parameters used to augment the F-Actin dataset using `pySTED`. For each acquisition, a set of parameter was uniformly sampled from the provided range.

| Models | Parameters | Fluorophore A | Fluorophore B |
|---|---|---|---|
| LinTSDiag | Exc. | 1.9 µW | 1.7 µW |
| | STED | 249 mW | 238 mW |
| | Pdt. | 19 µs | 17 µs |
| Kernel-TS | Exc. | 1.6 µW | 1.8 µW |
| | STED | 300 mW | 240 mW |
| | Pdt. | 21 µs | 21 µs |

Supplementary Tab. 9: Parameters at convergence of LinTSDiag and Kernel-TS for the two fluorophores in simulation. These parameters were used to acquire the images in Figure 3d.

| Specifications | Parameters | HS+LP | HSLP | HSHP | LSLP | LSHP |
|---|---|---|---|---|---|---|
| Photobleaching | Exc. | 10 µW | 10 µW | 2 µW | 10 µW | 2 µW |
| | STED | 150 mW | 150 mW | 150 mW | 150 mW | 150 mW |
| | Pdt. | 30 µs | 30 µs | 25 µs | 30 µs | 25 µs |
| | Target | 0.2 | 0.2 | 0.7 | 0.2 | 0.7 |
| Signal | Exc. | 1 µW | 10 µW | 10 µW | 10 µW | 10 µW |
| | STED | 0 W | 0 W | 0 W | 0 W | 0 W |
| | Pdt. | 10 µs | 10 µs | 10 µs | 10 µs | 10 µs |
| | Target | 200 | 200 | 200 | 30 | 30 |

Supplementary Tab. 10: Imaging parameters and target values to generate the synthetic fluorophores properties that are used for the evaluation of the RL model. HS+LP: High-Signal+ Low-Photobleaching, HSLP: High-Signal Low-Photobleaching, HSHP: High-Signal High-Photobleaching, Low-Signal Low-Photobleaching and LSHP: Low-Signal High-Photobleaching

| Dataset | Images |
|---|---|
| Actin [3] | 1224 |
| Tubulin [3] | 309 |
| PSD95 [3] | 246 |
| $\alpha$CaMKII [3] | 280 |
| LifeAct [3] | 246 |
| PSD95-Bassoon [4] | 3549 |
| Total | 5854 |

Supplementary Tab. 11: Datasets used to train the RL model. Images with a quality rating $> 0.7$ from Durand *et al.* [3], manually obtained from an expert, were kept for training. 2-color images of PSD95 and Bassoon from Wiesner *et al.* [4] were used. Only the Bassoon channel was kept for training. Full images were cropped to $224 \times 224$ pixels with an overlap of 25%. Crops containing at least 10% of foreground were kept. The foreground mask was obtained by a Gaussian filtered image ($sigma = 5\,\text{px}$) from the sum of both channels (PSD95 and Bassoon) followed by the Triangle method.

| Specifications | Parameters | Range of values |
|---|---|---|
| Photobleaching | Exc. | [1, 10] µW |
| | STED | [100, 200] mW |
| | Pdt. | [5, 75] µs |
| | Target | [0.1, 0.9] |
| Signal | Exc. | [1, 10] µW |
| | STED | 0 W |
| | Pdt. | 10 µs |
| | Target | [10, 400] |

Supplementary Tab. 12: Properties used to randomly generate fluorophores on-the-fly. Values are uniformly sampled and the fluorophore properties $k_1$, $b$ and $\sigma_{\text{abs}}$ are optimized for their sampled target.

| Structure | Value |
|---|---|
| CaMKII-$\beta$ | 2.25 |
| PSD95 | 2.25 |
| Bassoon | 2.25 |
| F-Actin | 3.0 |
| Tubulin | 3.75 |
| Lifeact | 3.75 |

Supplementary Tab. 13: Scaling factor used during the optimization process of the fluorophore properties. These scaling factors are used to ensure a proper number of detected photons. Denser structure require a bigger scaling factor. These factors were determined empirically.

| Parameter | Value |
|---|---|
| Figure 3b-d | |
| STED power | [0, 300] mW |
| Excitation power | [0, 2] µW |
| Pixel dwelltime | [0, 30] µs |
| Extended Fig 2 | |
| STED power | [0, 300] mW |
| Excitation power | [0, 2] µW |
| Pixel dwelltime | [0, 30] µs |
| Figure 3e-f | |
| STED power | [0, 50] % |
| Excitation power | [0, 3] % |
| Pixel dwelltime | [1, 30] µs |
| Linesteps | [1, 5] |
| Figure 3g-h | |
| STED power (STAR-ORANGE) | [0, 50] % |
| Excitation power (STAR-ORANGE) | [0, 20] % |
| Linesteps (STAR-ORANGE) | [1, 6] |
| STED power (STAR-RED) | [0, 50] % |
| Excitation power (STAR-RED) | [0, 4] % |
| Linesteps (STAR-RED) | [1, 6] |
| Figure 4c-g | |
| STED power | [0, 300] mW |
| Excitation power | [0, 2] µW |
| Pixel dwelltime | [0, 30] ms |
| Threshold 0 | [0, 15] |
| Threshold 1 | [0, 15] |
| Decision time 0 | [0, 5] µs |
| Decision time 1 | [0, 5] µs |
| Figure 4h-j | |
| STED power | [0, 50] % |
| Excitation power | [0, 15] % |
| Pixel dwelltime | [1, 30] µs |
| Threshold 0 | [1, 15] |
| Decision time 1 | [10, 40] % |
| Threshold 0 | [1, 15] |
| Decision time 1 | [10, 40] % |

Supplementary Tab. 14: Range of imaging parameters that were used for the bandit optimization.

| Parameter | Value |
|---|---|
| Figure 5c-f and Supplementary Fig 10 | |
| STED power | [0, 350] mW |
| Excitation power | [0, 20] µW |
| Pixel dwelltime | [1, 100] µs |
| Figure 6a-c and Extended Fig. 3a-b | |
| STED power | [0, 80] % |
| Excitation power | [0, 18] % |
| Pixel dwelltime | [1, 100] µs |
| Figure 6d and Extended Fig. 4 | |
| STED power | [0, 50] % |
| Excitation power | [0, 3] % |
| Pixel dwelltime | [1, 60] µs |
| Figure 6e | |
| STED power | [0, 50] % |
| Excitation power | [0, 10] % |
| Pixel dwelltime | [1, 60] µs |

Supplementary Tab. 15: Range of imaging parameters that were used for the RL optimization.

| | Species | Supplier | Category | Dilution | Related Fig. |
|---|---|---|---|---|---|
| PSD95 | Mouse | Invitrogen | MA1-045 | 1:500 | 4h-j, 6b |
| CaMKII-beta | Mouse | Thermo Fish. | 139800 | 1:50 | 6a |
| Tom20 | Rabbit | Santa Cruz | sc-11415 (FL-145) | 1:400 | 6c |
| Bassoon | Rabbit | Synaptic Systems | 141003 | 1:500 | S 3-5 |

Supplementary Tab. 16: Primary antibodies Table

| Fluorophores | Host | Reactivity | Supplier | Category | Dilution | Related Fig. |
|---|---|---|---|---|---|---|
| STAR635P | Goat | Mouse | Abberior | ST635P-1002-500UG | 1:250 | 4h-j 6a-c |
| STAR635P | Goat | Rabbit | Abberior | ST635P-1001-500UG | 1:250 | 6c |
| ATTO647N | Goat | Rabbit | Rockland | 611-156-122 | 1:250 | S 3-5 |

Supplementary Tab. 17: Secondary antibodies Table

| idx | Layer | Kernel size | Input size | Output size | Num. parameters |
|---|---|---|---|---|---|
| 0 | UNet (UNet) | – | [1, 1, 224, 224] | [1, 1, 224, 224] | – |
| 1 | -DoubleConv (inc): 1-1 | – | [1, 1, 224, 224] | [1, 64, 224, 224] | – |
| 2 | - -Sequential (double_conv): 2-1 | – | [1, 1, 224, 224] | [1, 64, 224, 224] | – |
| 3 | - - -Conv2d (0): 3-1 | [3, 3] | [1, 1, 224, 224] | [1, 64, 224, 224] | 576 |
| 4 | - - -BatchNorm2d (1): 3-2 | – | [1, 64, 224, 224] | [1, 64, 224, 224] | 128 |
| 5 | - - -ReLU (2): 3-3 | – | [1, 64, 224, 224] | [1, 64, 224, 224] | – |
| 6 | - - -Conv2d (3): 3-4 | [3, 3] | [1, 64, 224, 224] | [1, 64, 224, 224] | 36,864 |
| 7 | - - -BatchNorm2d (4): 3-5 | – | [1, 64, 224, 224] | [1, 64, 224, 224] | 128 |
| 8 | - - -ReLU (5): 3-6 | – | [1, 64, 224, 224] | [1, 64, 224, 224] | – |
| 9 | -Down (down1): 1-2 | – | [1, 64, 224, 224] | [1, 128, 112, 112] | – |
| 10 | - -Sequential (maxpool_conv): 2-2 | – | [1, 64, 224, 224] | [1, 128, 112, 112] | – |
| 11 | - - -MaxPool2d (0): 3-7 | 2 | [1, 64, 224, 224] | [1, 64, 112, 112] | – |
| 12 | - - -DoubleConv (1): 3-8 | – | [1, 64, 112, 112] | [1, 128, 112, 112] | – |
| 13 | - - - -Sequential (double_conv): 4-1 | – | [1, 64, 112, 112] | [1, 128, 112, 112] | – |
| 14 | - - - - -Conv2d (0): 5-1 | [3, 3] | [1, 64, 112, 112] | [1, 128, 112, 112] | 73,728 |
| 15 | - - - - -BatchNorm2d (1): 5-2 | – | [1, 128, 112, 112] | [1, 128, 112, 112] | 256 |
| 16 | - - - - -ReLU (2): 5-3 | – | [1, 128, 112, 112] | [1, 128, 112, 112] | – |
| 17 | - - - - -Conv2d (3): 5-4 | [3, 3] | [1, 128, 112, 112] | [1, 128, 112, 112] | 147,456 |
| 18 | - - - - -BatchNorm2d (4): 5-5 | – | [1, 128, 112, 112] | [1, 128, 112, 112] | 256 |
| 19 | - - - - -ReLU (5): 5-6 | – | [1, 128, 112, 112] | [1, 128, 112, 112] | – |
| 20 | -Down (down2): 1-3 | – | [1, 128, 112, 112] | [1, 256, 56, 56] | – |
| 21 | - -Sequential (maxpool_conv): 2-3 | – | [1, 128, 112, 112] | [1, 256, 56, 56] | – |
| 22 | - - -MaxPool2d (0): 3-9 | 2 | [1, 128, 112, 112] | [1, 128, 56, 56] | – |
| 23 | - - -DoubleConv (1): 3-10 | – | [1, 128, 56, 56] | [1, 256, 56, 56] | – |
| 24 | - - - -Sequential (double_conv): 4-2 | – | [1, 128, 56, 56] | [1, 256, 56, 56] | – |
| 25 | - - - - -Conv2d (0): 5-7 | [3, 3] | [1, 128, 56, 56] | [1, 256, 56, 56] | 294,912 |
| 26 | - - - - -BatchNorm2d (1): 5-8 | – | [1, 256, 56, 56] | [1, 256, 56, 56] | 512 |
| 27 | - - - - -ReLU (2): 5-9 | – | [1, 256, 56, 56] | [1, 256, 56, 56] | – |
| 28 | - - - - -Conv2d (3): 5-10 | [3, 3] | [1, 256, 56, 56] | [1, 256, 56, 56] | 589,824 |
| 29 | - - - - -BatchNorm2d (4): 5-11 | – | [1, 256, 56, 56] | [1, 256, 56, 56] | 512 |
| 30 | - - - - -ReLU (5): 5-12 | – | [1, 256, 56, 56] | [1, 256, 56, 56] | – |
| 31 | -Down (down3): 1-4 | – | [1, 256, 56, 56] | [1, 512, 28, 28] | – |
| 32 | - -Sequential (maxpool_conv): 2-4 | – | [1, 256, 56, 56] | [1, 512, 28, 28] | – |
| 33 | - - -MaxPool2d (0): 3-11 | 2 | [1, 256, 56, 56] | [1, 256, 28, 28] | – |
| 34 | - - -DoubleConv (1): 3-12 | – | [1, 256, 28, 28] | [1, 512, 28, 28] | – |
| 35 | - - - -Sequential (double_conv): 4-3 | – | [1, 256, 28, 28] | [1, 512, 28, 28] | – |
| 36 | - - - - -Conv2d (0): 5-13 | [3, 3] | [1, 256, 28, 28] | [1, 512, 28, 28] | 1,179,648 |
| 37 | - - - - -BatchNorm2d (1): 5-14 | – | [1, 512, 28, 28] | [1, 512, 28, 28] | 1,024 |
| 38 | - - - - -ReLU (2): 5-15 | – | [1, 512, 28, 28] | [1, 512, 28, 28] | – |
| 39 | - - - - -Conv2d (3): 5-16 | [3, 3] | [1, 512, 28, 28] | [1, 512, 28, 28] | 2,359,296 |
| 40 | - - - - -BatchNorm2d (4): 5-17 | – | [1, 512, 28, 28] | [1, 512, 28, 28] | 1,024 |
| 41 | - - - - -ReLU (5): 5-18 | – | [1, 512, 28, 28] | [1, 512, 28, 28] | – |
| 42 | -Down (down4): 1-5 | – | [1, 512, 28, 28] | [1, 1024, 14, 14] | – |
| 43 | - -Sequential (maxpool_conv): 2-5 | – | [1, 512, 28, 28] | [1, 1024, 14, 14] | – |
| 44 | - - -MaxPool2d (0): 3-13 | 2 | [1, 512, 28, 28] | [1, 512, 14, 14] | – |
| 45 | - - -DoubleConv (1): 3-14 | – | [1, 512, 14, 14] | [1, 1024, 14, 14] | – |
| 46 | - - - -Sequential (double_conv): 4-4 | – | [1, 512, 14, 14] | [1, 1024, 14, 14] | – |
| 47 | - - - - -Conv2d (0): 5-19 | [3, 3] | [1, 512, 14, 14] | [1, 1024, 14, 14] | 4,718,592 |
| 48 | - - - - -BatchNorm2d (1): 5-20 | – | [1, 1024, 14, 14] | [1, 1024, 14, 14] | 2,048 |
| 49 | - - - - -ReLU (2): 5-21 | – | [1, 1024, 14, 14] | [1, 1024, 14, 14] | – |
| 50 | - - - - -Conv2d (3): 5-22 | [3, 3] | [1, 1024, 14, 14] | [1, 1024, 14, 14] | 9,437,184 |
| 51 | - - - - -BatchNorm2d (4): 5-23 | – | [1, 1024, 14, 14] | [1, 1024, 14, 14] | 2,048 |
| 52 | - - - - -ReLU (5): 5-24 | – | [1, 1024, 14, 14] | [1, 1024, 14, 14] | – |
| 53 | -Up (up1): 1-6 | – | [1, 1024, 14, 14] | [1, 512, 28, 28] | – |
| 54 | - -ConvTranspose2d (up): 2-6 | [2, 2] | [1, 1024, 14, 14] | [1, 512, 28, 28] | 2,097,664 |
| 55 | - -DoubleConv (conv): 2-7 | – | [1, 1024, 28, 28] | [1, 512, 28, 28] | – |
| 56 | - - -Sequential (double_conv): 3-15 | – | [1, 1024, 28, 28] | [1, 512, 28, 28] | – |
| 57 | - - - -Conv2d (0): 4-5 | [3, 3] | [1, 1024, 28, 28] | [1, 512, 28, 28] | 4,718,592 |
| 58 | - - - -BatchNorm2d (1): 4-6 | – | [1, 512, 28, 28] | [1, 512, 28, 28] | 1,024 |
| 59 | - - - -ReLU (2): 4-7 | – | [1, 512, 28, 28] | [1, 512, 28, 28] | – |
| 60 | - - - -Conv2d (3): 4-8 | [3, 3] | [1, 512, 28, 28] | [1, 512, 28, 28] | 2,359,296 |
| 61 | - - - -BatchNorm2d (4): 4-9 | – | [1, 512, 28, 28] | [1, 512, 28, 28] | 1,024 |
| 62 | - - - -ReLU (5): 4-10 | – | [1, 512, 28, 28] | [1, 512, 28, 28] | – |
| 63 | -Up (up2): 1-7 | – | [1, 512, 28, 28] | [1, 256, 56, 56] | – |
| 64 | - -ConvTranspose2d (up): 2-8 | [2, 2] | [1, 512, 28, 28] | [1, 256, 56, 56] | 524,544 |
| 65 | - -DoubleConv (conv): 2-9 | – | [1, 512, 56, 56] | [1, 256, 56, 56] | – |
| 66 | - - -Sequential (double_conv): 3-16 | – | [1, 512, 56, 56] | [1, 256, 56, 56] | – |
| 67 | - - - -Conv2d (0): 4-11 | [3, 3] | [1, 512, 56, 56] | [1, 256, 56, 56] | 1,179,648 |
| 68 | - - - -BatchNorm2d (1): 4-12 | – | [1, 256, 56, 56] | [1, 256, 56, 56] | 512 |
| 69 | - - - -ReLU (2): 4-13 | – | [1, 256, 56, 56] | [1, 256, 56, 56] | – |
| 70 | - - - -Conv2d (3): 4-14 | [3, 3] | [1, 256, 56, 56] | [1, 256, 56, 56] | 589,824 |
| 71 | - - - -BatchNorm2d (4): 4-15 | – | [1, 256, 56, 56] | [1, 256, 56, 56] | 512 |
| 72 | - - - -ReLU (5): 4-16 | – | [1, 256, 56, 56] | [1, 256, 56, 56] | – |
| 73 | -Up (up3): 1-8 | – | [1, 256, 56, 56] | [1, 128, 112, 112] | – |
| 74 | - -ConvTranspose2d (up): 2-10 | [2, 2] | [1, 256, 56, 56] | [1, 128, 112, 112] | 131,200 |
| 75 | - -DoubleConv (conv): 2-11 | – | [1, 256, 112, 112] | [1, 128, 112, 112] | – |
| 76 | - - -Sequential (double_conv): 3-17 | – | [1, 256, 112, 112] | [1, 128, 112, 112] | – |
| 77 | - - - -Conv2d (0): 4-17 | [3, 3] | [1, 256, 112, 112] | [1, 128, 112, 112] | 294,912 |
| 78 | - - - -BatchNorm2d (1): 4-18 | – | [1, 128, 112, 112] | [1, 128, 112, 112] | 256 |
| 79 | - - - -ReLU (2): 4-19 | – | [1, 128, 112, 112] | [1, 128, 112, 112] | – |
| 80 | - - - -Conv2d (3): 4-20 | [3, 3] | [1, 128, 112, 112] | [1, 128, 112, 112] | 147,456 |
| 81 | - - - -BatchNorm2d (4): 4-21 | – | [1, 128, 112, 112] | [1, 128, 112, 112] | 256 |
| 82 | - - - -ReLU (5): 4-22 | – | [1, 128, 112, 112] | [1, 128, 112, 112] | – |
| 83 | -Up (up4): 1-9 | – | [1, 128, 112, 112] | [1, 64, 224, 224] | – |
| 84 | - -ConvTranspose2d (up): 2-12 | [2, 2] | [1, 128, 112, 112] | [1, 64, 224, 224] | 32,832 |
| 85 | - -DoubleConv (conv): 2-13 | – | [1, 128, 224, 224] | [1, 64, 224, 224] | – |
| 86 | - - -Sequential (double_conv): 3-18 | – | [1, 128, 224, 224] | [1, 64, 224, 224] | – |
| 87 | - - - -Conv2d (0): 4-23 | [3, 3] | [1, 128, 224, 224] | [1, 64, 224, 224] | 73,728 |
| 88 | - - - -BatchNorm2d (1): 4-24 | – | [1, 64, 224, 224] | [1, 64, 224, 224] | 128 |
| 89 | - - - -ReLU (2): 4-25 | – | [1, 64, 224, 224] | [1, 64, 224, 224] | – |
| 90 | - - - -Conv2d (3): 4-26 | [3, 3] | [1, 64, 224, 224] | [1, 64, 224, 224] | 36,864 |
| 91 | - - - -BatchNorm2d (4): 4-27 | – | [1, 64, 224, 224] | [1, 64, 224, 224] | 128 |
| 92 | - - - -ReLU (5): 4-28 | – | [1, 64, 224, 224] | [1, 64, 224, 224] | – |
| 93 | -OutConv (outc): 1-10 | – | [1, 64, 224, 224] | [1, 1, 224, 224] | – |
| 94 | - -Conv2d (conv): 2-14 | [1, 1] | [1, 64, 224, 224] | [1, 1, 224, 224] | 65 |
| 95 | -Sigmoid (sigmoid): 1-11 | – | [1, 1, 224, 224] | [1, 1, 224, 224] | – |

Supplementary Tab. 18: U-Net$_{\text{datamap}}$ architecture.

| idx | Layer | Kernel size | Input size | Output size | Num. parameters |
|---|---|---|---|---|---|
| 0 | LinearModel (LinearModel) | – | [1, 3] | [1, 1] | – |
| 1 | -Sequential (feature_extractor): 1-1 | – | [1, 3] | [1, 32] | – |
| 2 | - -Linear (0): 2-1 | – | [1, 3] | [1, 32] | 128 |
| 3 | - -ReLU (1): 2-2 | – | [1, 32] | [1, 32] | – |
| 4 | - -Dropout (2): 2-3 | – | [1, 32] | [1, 32] | – |
| 5 | - -Linear (3): 2-4 | – | [1, 32] | [1, 32] | 1,056 |
| 6 | - -ReLU (4): 2-5 | – | [1, 32] | [1, 32] | – |
| 7 | - -Dropout (5): 2-6 | – | [1, 32] | [1, 32] | – |
| 8 | -Linear (linear): 1-2 | – | [1, 32] | [1, 1] | 32 |

Supplementary Tab. 19: LinTSDiag model without contextual information

| idx | Layer | Kernel size | Input size | Output size | Num. parameters |
|---|---|---|---|---|---|
| 0 | ImageContextLinearModel (ImageContextLinearModel) | – | [1, 7] | [1, 1, 1] | – |
| 1 | -Sequential (feature_extractor): 1-1 | – | [1, 1, 7] | [1, 1, 32] | – |
| 2 | - -Linear (0): 2-1 | – | [1, 1, 7] | [1, 1, 32] | 256 |
| 3 | - -ReLU (1): 2-2 | – | [1, 1, 32] | [1, 1, 32] | – |
| 4 | - -Dropout (2): 2-3 | – | [1, 1, 32] | [1, 1, 32] | – |
| 5 | - -Linear (3): 2-4 | – | [1, 1, 32] | [1, 1, 32] | 1,056 |
| 6 | - -ReLU (4): 2-5 | – | [1, 1, 32] | [1, 1, 32] | – |
| 7 | - -Dropout (5): 2-6 | – | [1, 1, 32] | [1, 1, 32] | – |
| 8 | -ContextEncoder (context_encoder): 1-2 | – | [1, 1, 224, 224] | [1, 32] | – |
| 9 | - -Sequential (context_encoder): 2-7 | – | [1, 1, 224, 224] | [1, 32] | – |
| 10 | - - -Conv2d (0): 3-1 | [3, 3] | [1, 1, 224, 224] | [1, 8, 224, 224] | 80 |
| 11 | - - -BatchNorm2d (1): 3-2 | – | [1, 8, 224, 224] | [1, 8, 224, 224] | 16 |
| 12 | - - -MaxPool2d (2): 3-3 | 4 | [1, 8, 224, 224] | [1, 8, 56, 56] | – |
| 13 | - - -ReLU (3): 3-4 | – | [1, 8, 56, 56] | [1, 8, 56, 56] | – |
| 14 | - - -Conv2d (4): 3-5 | [3, 3] | [1, 8, 56, 56] | [1, 16, 56, 56] | 1,168 |
| 15 | - - -BatchNorm2d (5): 3-6 | – | [1, 16, 56, 56] | [1, 16, 56, 56] | 32 |
| 16 | - - -AdaptiveAvgPool2d (6): 3-7 | – | [1, 16, 56, 56] | [1, 16, 1, 1] | – |
| 17 | - - -ReLU (7): 3-8 | – | [1, 16, 1, 1] | [1, 16, 1, 1] | – |
| 18 | - - -Flatten (8): 3-9 | – | [1, 16, 1, 1] | [1, 16] | – |
| 19 | - - -Dropout (9): 3-10 | – | [1, 16] | [1, 16] | – |
| 20 | - - -Linear (10): 3-11 | – | [1, 16] | [1, 32] | 544 |
| 21 | - - -ReLU (11): 3-12 | – | [1, 32] | [1, 32] | – |
| 22 | - - -Dropout (12): 3-13 | – | [1, 32] | [1, 32] | – |
| 23 | -Linear (pre): 1-3 | – | [1, 1, 64] | [1, 1, 64] | 4,160 |
| 24 | -Linear (linear): 1-4 | – | [1, 1, 64] | [1, 1, 1] | 64 |

Supplementary Tab. 20: LinTSDiag model with contextual information. The output from layer 7 and 22 are concatenated and given as input to layer 23.

| idx | Layer | Kernel size | Input size | Output size | Num. parameters |
|---|---|---|---|---|---|
| 0 | PooledPolicy (PooledPolicy) | – | [1, 3, 224, 224] | [1, 3] | 3 |
| 1 | -ModuleList (image_encoder_layers): 1-1 | – | – | – | – |
| 2 | - -Conv2d (0): 2-1 | [8, 8] | [1, 3, 224, 224] | [1, 32, 55, 55] | 6,176 |
| 3 | - -Conv2d (1): 2-2 | [4, 4] | [1, 32, 55, 55] | [1, 64, 26, 26] | 32,832 |
| 4 | - -Conv2d (2): 2-3 | [3, 3] | [1, 64, 26, 26] | [1, 64, 24, 24] | 36,928 |
| 5 | -AdaptiveMaxPool2d (image_pool_layer): 1-2 | – | [1, 64, 24, 24] | [1, 64, 1, 1] | – |
| 6 | -ModuleList (signal_encoder_layers): 1-3 | – | – | – | – |
| 7 | - -Linear (0): 2-4 | – | [1, 181] | [1, 16] | 2,912 |
| 8 | - -Linear (1): 2-5 | – | [1, 16] | [1, 16] | 272 |
| 9 | -Linear (policy_to_actions_layer): 1-4 | – | [1, 80] | [1, 3] | 243 |

Supplementary Tab. 21: Policy model used for the RL experiments.

| idx | Layer | Kernel size | Input size | Output size | Num. parameters |
|---|---|---|---|---|---|
| 0 | PooledValueFunction (PooledValueFunction) | – | [1, 3, 224, 224] | [1, 1] | – |
| 1 | -ModuleList (image_encoder_layers): 1-1 | – | – | – | – |
| 2 | - -Conv2d (0): 2-1 | [8, 8] | [1, 3, 224, 224] | [1, 32, 55, 55] | 6,176 |
| 3 | - -Conv2d (1): 2-2 | [4, 4] | [1, 32, 55, 55] | [1, 64, 26, 26] | 32,832 |
| 4 | - -Conv2d (2): 2-3 | [3, 3] | [1, 64, 26, 26] | [1, 64, 24, 24] | 36,928 |
| 5 | -AdaptiveMaxPool2d (image_pool_layer): 1-2 | – | [1, 64, 24, 24] | [1, 64, 1, 1] | – |
| 6 | -ModuleList (signal_encoder_layers): 1-3 | – | – | – | – |
| 7 | - -Linear (0): 2-4 | – | [1, 181] | [1, 16] | 2,912 |
| 8 | - -Linear (1): 2-5 | – | [1, 16] | [1, 16] | 272 |
| 9 | -Linear (linear1): 1-4 | – | [1, 80] | [1, 64] | 5,184 |
| 10 | -Linear (linear2): 1-5 | – | [1, 64] | [1, 1] | 65 |

Supplementary Tab. 22: Value function used for the RL experiments.

### Figures

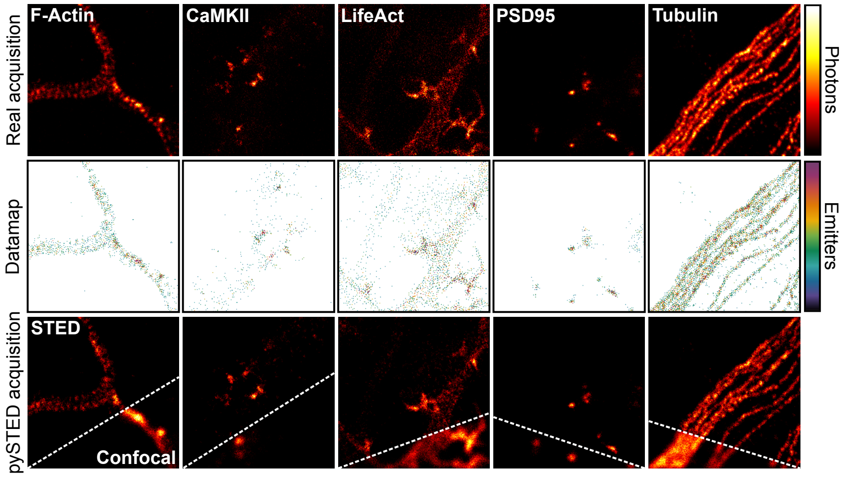


Supplementary Fig. 1: Examples of datamap generation using the trained U-Net$_{\text{datamap}}$ model. A simulated `pySTED` acquisition of a STED and confocal image is shown. All images are normalized using min-max.

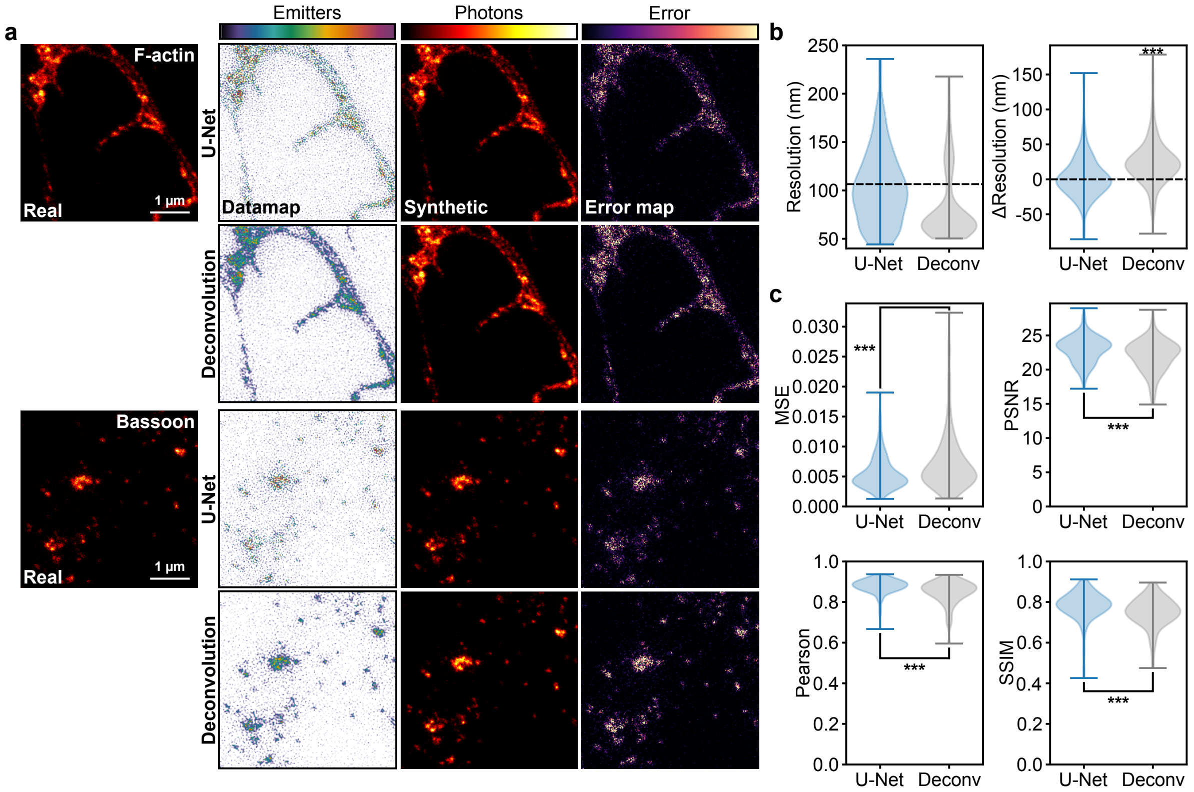


Supplementary Fig. 2: Datamap generation using the trained U-Net$_{\text{datamap}}$ model or a Richardson-Lucy deconvolution. a) Two representative examples from an independent image set are displayed (top: F-actin; bottom: Bassoon). (left) Datamap generated by the U-Net$_{\text{datamap}}$ or Richardson-Lucy deconvolution. (center) A synthetic image is generated in `pySTED` from the datamap. (right) The pixel-wise error map between the real and synthetic images. b) (left) The resolution of the synthetic images generated using datamaps from the U-Net$_{\text{datamap}}$ or the Richardson-Lucy deconvolution (Deconv). Black dashed line is the average resolution measured on real images. (right) The resolution of images resulting from datamaps generated by the U-Net$_{\text{datamap}}$ is not significantly different from the one of real images ($p$-value=0.5858; Wilcoxon signed-rank test) but deconvolution leads in an overestimation of the resolution ($p$-value=$9.4973 \times 10^{-20}$, Wilcoxon signed-rank test). Black dashed line is at zero value. c) Quantitative measurements comparing the synthetic images and their real version (MSE: mean squared error; PSNR: peak signal to noise ration; Pearson: Pearson correlation coefficient; SSIM: structural similarity index measure). Statistical analysis reveals that images generated with datamaps from U-Net$_{\text{datamap}}$ are more similar to the real version compared to images generated from datamaps obtained from Richardson-Lucy deconvolution ($p$-value$_{\text{MSE}}$=$4.4129 \times 10^{-21}$, $p$-value$_{\text{PSNR}}$=$1.1484 \times 10^{-22}$, $p$-value$_{\text{Pearson}}$=$9.7045 \times 10^{-31}$, and $p$-value$_{\text{SSIM}}$=$3.7712 \times 10^{-26}$; Wilcoxon signed-rank test). N=180 images.

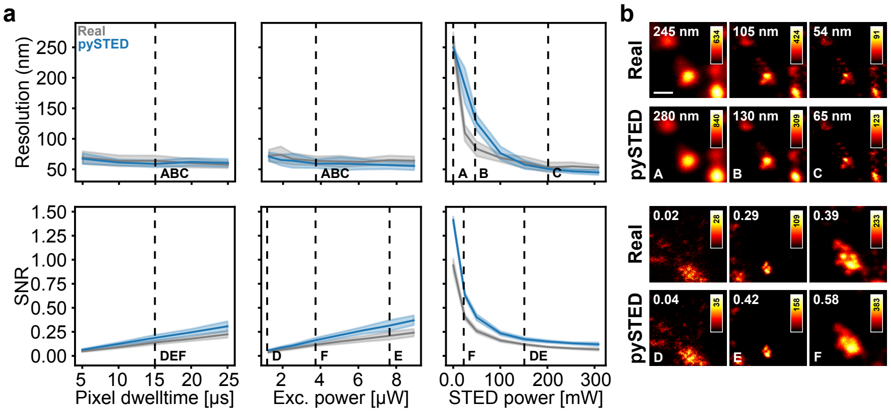


Supplementary Fig. 3: Comparison of the pySTED simulations with acquisitions on a STED microscope. a) The pixel dwelltime, excitation and depletion laser power were varied in the pySTED simulation environment (blue) and compared with an equivalent range on the microscope (grey) (Default parameters; Pixel dwelltime: 15 µs, Excitation power: 3.76 µW, STED power: 150 mW). Depicted here is the mean and 1 standard deviation (shade) obtained from 10 acquisitions of each parameter combinations for two imaging optimization objectives : resolution and signal ratio (SNR). For relatively low STED laser power ([0, 100] mW), pySTED underestimates the resolution of the image. For the signal ratio, pySTED slightly overestimates but keeps similar variations as a function of the imaging parameters (bottom). Increasing the pixel dwelltime and the excitation power while keeping the other parameters to their default values leads in an increased signal ratio. Increasing the STED power reduces the signal ratio similarly to a real acquisition. This is expected from the presence of the depletion beam which depletes the fluorophores. The evolution of the resolution and signal ratio are shown for other parameters that can be modulated in pySTED such as gating delay, gating time, and the number of line repetitions in Supplementary Fig. 5. b) Images acquired on a real microscope and their corresponding pySTED simulation for different combinations of imaging parameters (bottom left letters; see vertical dashed black lines in a). The measured imaging optimization objectives are shown on the top left of each image (top row: resolution, bottom row: SNR). Scale bar: 500 nm.

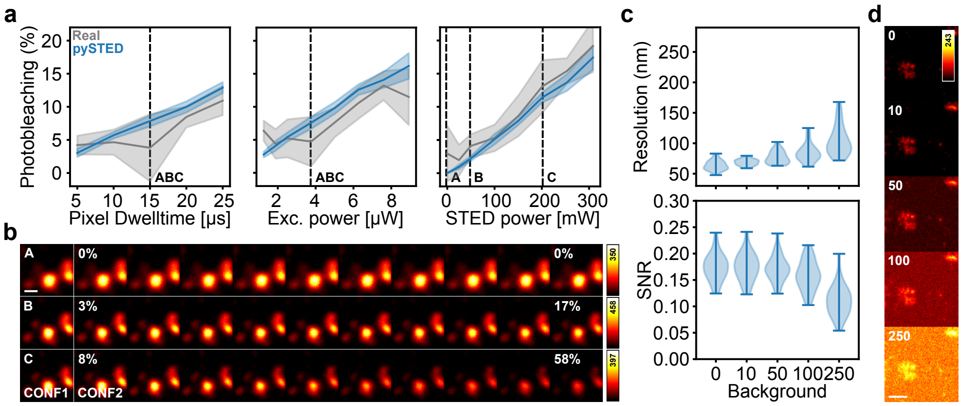


Supplementary Fig. 4: Validation of the fluorophore properties of pySTED with a real sample. a) The imaging parameters were varied across the range of parameters in pySTED (blue) and on a real microscope (gray) by keeping other parameters constant (Default parameters: Pixel dwelltime: 15 µs, Excitation power: 3.76 µW, STED power: 150 mW). The mean (solid line) and standard deviation (shade) of 10 repetitions are presented. b) Example of a sequential acquisition of 10 STED images with default parameters in pySTED. The first confocal is presented on the left to compare the signal loss in subsequent acquisition. After each STED acquisition, a confocal is acquired (CONF2) which allows to measure the photobleaching with respect to the original confocal (CONF1, Methods). The measured photobleaching is displayed on the top left of the images. The parameters selected for each line are displayed by vertical black dashed lines in a. c) pySTED allows to add background signal during the acquisition which impacts negatively the measured resolution and SNR. The distribution from 30 acquisitions is shown. Increasing the background signal in pySTED with constant acquisition parameters leads to an underestimation of the resolution and negatively impacts the measured signal ratio of the images d) Examples of acquisition with different background level displayed in c.

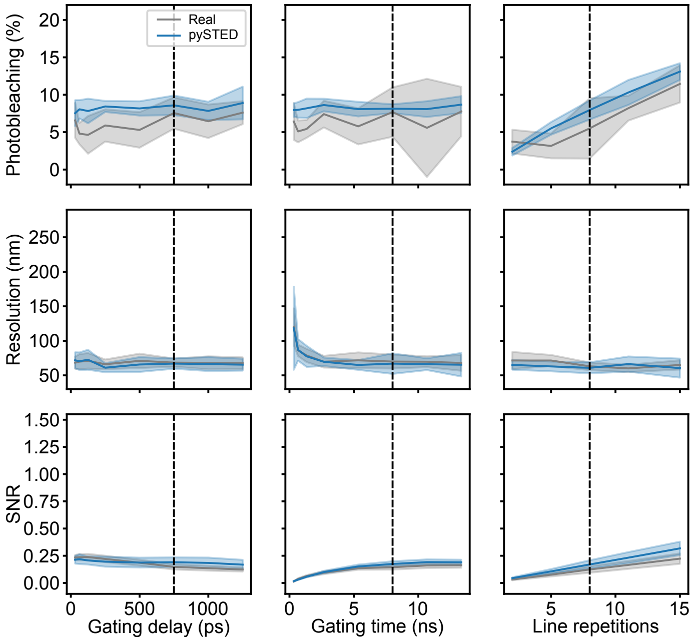


Supplementary Fig. 5: Evolution of the imaging optimization objectives (photobleaching, resolution and SNR) for the gating delay, gating time and the number of line repetitions. The pySTED acquisition (blue) is compared with the same parameters acquired on a real STED microscope (gray). Displayed is the mean (solid line) and 1 standard deviation (shade) from 10 repetitions.

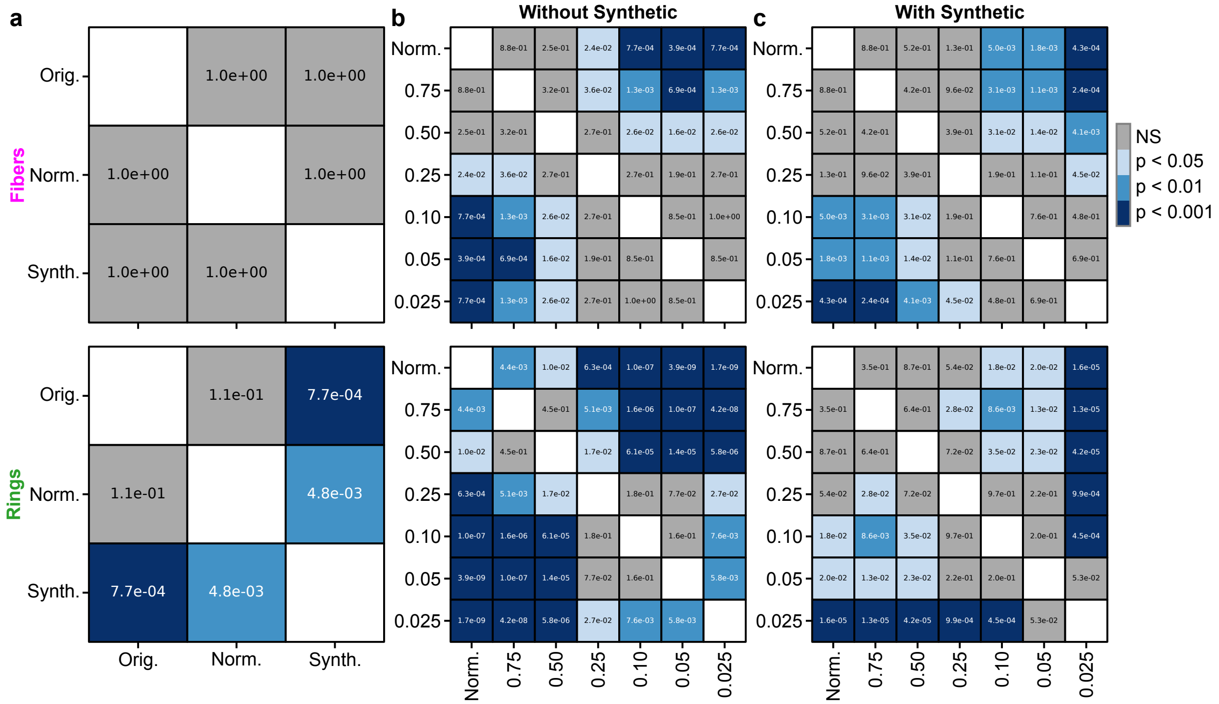


Supplementary Fig. 6: Statistical analysis of the performance of the trained U-Net for F-actin fibers (top row) and rings (bottom row) for values presented in Figure 2c-e. a) Comparison between original training (Orig.), updated normalization (Norm,) and with using synthetics images (Synth.). b) Comparison between the updated normalization and keeping a subset of images during training. c) Comparison between the updated normalization and keeping a subset of images that are augmented with pySTED.

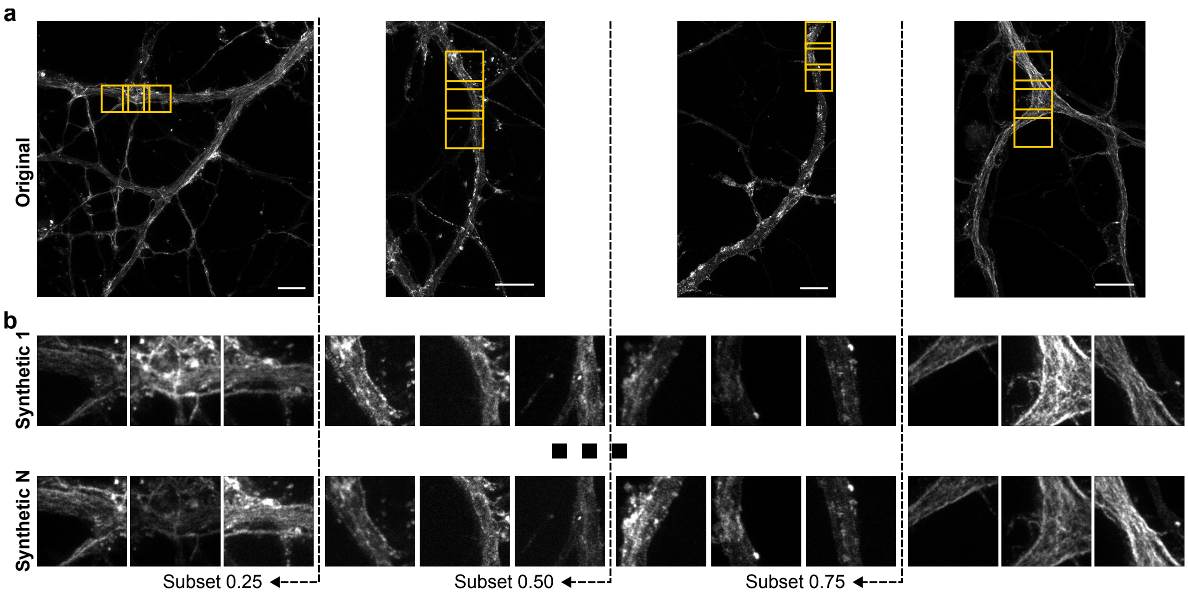


Supplementary Fig. 7: Schematic representation of training with subsets of images. a) In each subset, we remove the complete image and its respective valid crops to mimic how the annotation is typically performed, *i.e.* at the image level. Yellow overlays represent some of the valid crops that can be used for training. Not all crops are displayed. b) When synthetic images are added in the dataset, we augment each valid crop $N$ times to generate a more diverse training dataset. When training on a subset of images we remove the image and its respective synthetic crops from training. Crop size: $256 \times 256$ pixels. Scale bars: 5 µm.

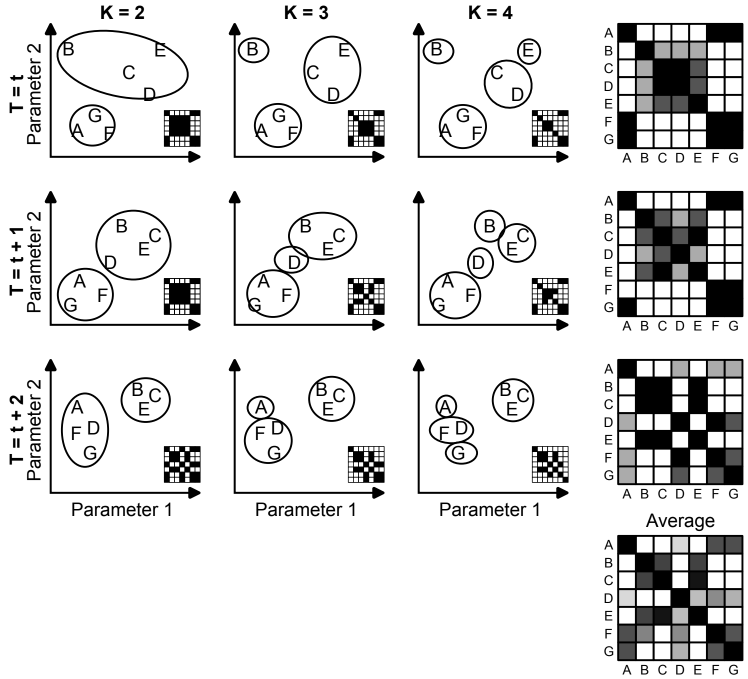


Supplementary Fig. 8: Schematic representation of the calculation of the correlation matrix from Figure 5c. At each time step $t$, $k$-means clustering is repeatedly computed with an increasing number of clusters resulting in an association matrix between each models. The average association matrix from $K$ clusters is then calculated (right column). To obtain the final correlation matrix, we average the association matrix from $t$ time steps (bottom right).

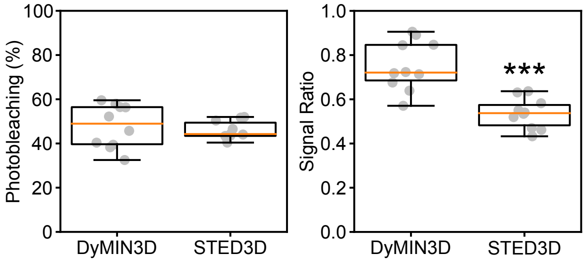


Supplementary Fig. 9: Comparison of the photobleaching and signal ratio between a DyMIN3D and STED3D acquisition using the same Excitation and STED power and pixel dwelltime. The photobleaching is not significantly different between the two conditions ($p = 0.5557$). The signal ratio is significantly increased for a DyMIN3D acquisition compared to conventional STED3D ($p = 6.2713 \times 10^{-5}$).

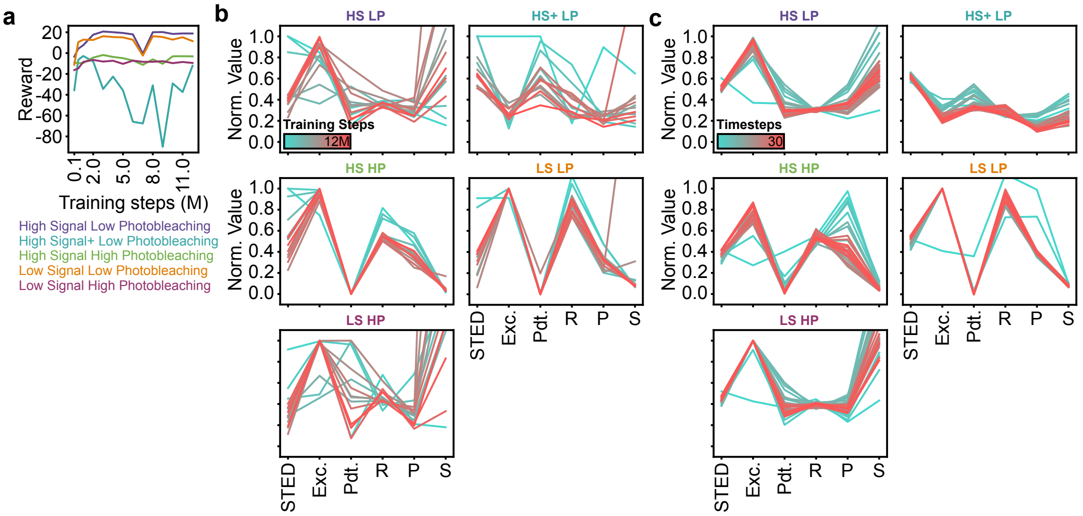


Supplementary Fig. 10: RL agent in simulation facing different fluorophore properties. a) Evolution of the average reward during an episode at the beginning (cyan, 100k timesteps) and at the end of training (red, 12M timesteps) for different fluorophore properties. b) Evolution of the policy (left) and imaging optimization objectives (right; R: Resolution, P: Photobleaching, S: Signal ratio) over the course of training (from cyan to red) for different fluorophore properties. c) Evolution of the policy (left) and imaging optimization objectives (right) after training (12M timesteps) during an episode for different fluorophore properties.

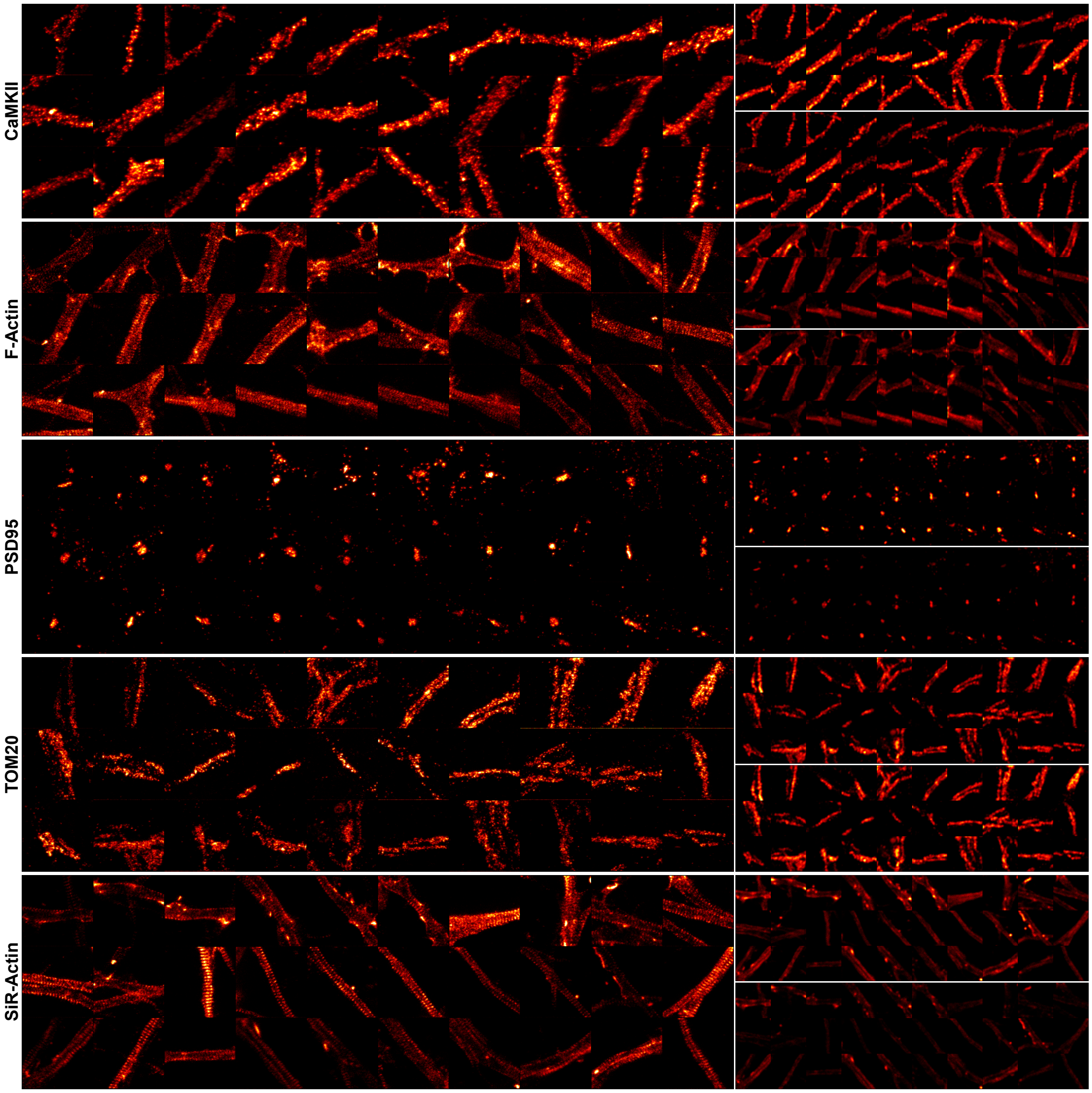


Supplementary Fig. 11: Images acquired by the RL agent in a real experiment. For each row, the sequence of acquired images goes from top left to bottom right. Right column: Confocal images before (top) and after (bottom) are presented for comparison. The confocal image after is normalized to the confocal before image. The STED images are normalized to the 99$^{th}$ percentile of the intensity of the confocal before image. Images are 5.12 µm × 5.12 µm.

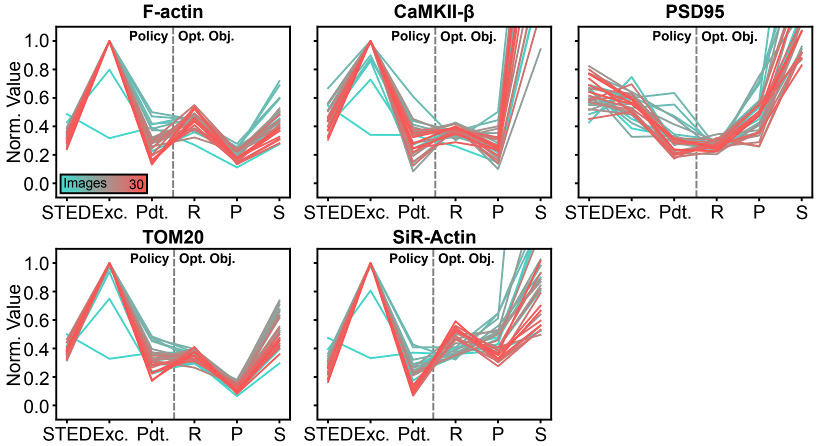


Supplementary Fig. 12: Evolution of the parameter selection (left; STED: STED power, Exc.: Excitation power, Pdt.: Pixel dwelltime) and imaging optimization objectives (right; R: Resolution, P: Photobleaching, S: Signal ratio) of the agent for the different structures.
